# CRISPR-targeted display of functional T cell receptors enables engineering of enhanced specificity and prediction of cross-reactivity

**DOI:** 10.1101/2020.06.23.166363

**Authors:** Rodrigo Vazquez-Lombardi, Johanna S. Jung, Florian Bieberich, Edo Kapetanovic, Erik Aznauryan, Cédric R. Weber, Sai T. Reddy

## Abstract

T cell receptor (TCR) gene therapy is a promising cell therapy approach for the treatment of cancer. However, most naturally occurring TCRs display low affinities to their peptide-MHC targets, and engineering of TCRs for enhanced affinity is complicated by the risk of introducing cross-reactivity and the poor correlation between affinity and function. Here we report the establishment of the TCR-accepting T cell (TnT) platform through five sequential CRISPR-Cas9 genome editing steps of a human T cell line, and demonstrate its application for functional engineering of TCRs and prediction of cross-reactivity. Using the TnT platform, we profile the mutational landscapes of tumor-specific TCRs at high-throughput to reveal a substantial discordance between antigen binding and antigen-induced signaling. Furthermore, we combine CRISPR-targeting, functional selection and deep sequencing to screen TCR mutagenesis libraries and identify variants with enhanced recognition of the cancer-testis antigen MAGE-A3. Finally, functional cross-reactivity profiling using TnT cells was able to accurately predict off-targets and identify engineered TCRs with exquisite specificity to MAGE-A3. Thus, the TnT platform represents a valuable technology for the engineering of TCRs with enhanced functional and safety profiles.

## INTRODUCTION

T cell receptor (TCR) gene therapy is a promising cell therapy approach against cancer that relies on viral transfer of genes encoding the α and β chains of tumor-reactive TCRs into autologous T cells, followed by their expansion and re-infusion into patients^1^. Unlike chimeric antigen receptor (CAR)-T cell therapies, which target surface tumor antigens, TCR-redirected T cells recognize processed intracellular tumor antigen peptides displayed by major histocompatibility complex (MHC) / human leukocyte antigen (HLA) molecules. This key feature vastly increases the number of potential antigen targets and is thought to allow for more efficient infiltration into solid tumors^2^, a known limitation of CAR-T cells^3^. As such, TCR gene therapy has been identified as an effective approach for driving durable responses against multiple cancers^2,4–12^. Despite its significant promise, the discovery, engineering and selection of optimal therapeutic TCRs remains a time-consuming process complicated by the low affinities of TCRs to their peptide-MHC targets and the inherent risk of TCR cross-reactivity^6,13,14^.

Different from antibodies, which typically recognize a single epitope, TCRs are able to recognize multiple peptide antigens presented by MHC^15^. For example, naturally-occurring TCRs have been shown to display a wide range of specificity profiles^16^, in some cases having the potential to recognize up to a million different peptides^17^. Therefore, cross-reactivity is vital to measure and subsequently avoid in order to engineer safe TCR gene therapies. In addition to specificity, TCR affinity and potency (i.e., functional avidity) are crucial parameters to consider when selecting optimal TCRs for gene therapy applications. Notably, TCR specificity, affinity and function are not necessarily correlated with each other. For example, while TCRs with high affinities to antigen (1-5 μM) tend to display high potency *in vitro*, TCRs with lower affinities (5-100 μM) often display a poor correlation between affinity and function^18,19^. In addition, engineered TCRs with supraphysiological affinities (< 1 μM) may display sub-optimal therapeutic activity due to increased T cell dysfunction^20,21^, inability to undergo serial TCR triggering^22^, and potential for reactivity against the presenting HLA molecule^23,24^.

Due to central and peripheral negative selection processes, naturally occurring high-affinity TCRs targeting self-antigens are extremely rare and their discovery requires extensive screening^25–27^. To overcome this, directed evolution and protein engineering methods that rely on display technologies (e.g., phage and yeast display) have been employed^28,29^. These require reformatting of TCRs into synthetic single-chain fragments, which are then screened on the surface of phage or yeast for binding towards peptide-MHC multimers. In one notable example, phage display was used to increase the affinity of a TCR targeting the cancer-testis antigen MAGE-A3. When applied as a TCR gene therapy, unexpected cross-reactivity towards a self-antigen expressed by beating cardiomyocytes cells was observed, which ultimately resulted in treatment-induced patient deaths in a clinical trial^14,30^. In light of this outcome, the development of TCR display platforms enabling simultaneous TCR engineering and detection of cross-reactivity on the basis of function (i.e., antigen-induced signaling) would be highly desirable. While a number of TCR engineering methods have been developed in mammalian cell lines^25,31–34^ and primary T cells^35,36^, these have only reported selections based on antigen binding. In addition, their use of viral transduction or plasmid transfection for TCR reconstitution is associated with limitations including random integration, constitutive expression and possible expression of different TCRs by a single cell.

Several technologies allowing the assessment of TCR specificity have been reported recently, highlighting a strong interest in the development and safety screening of TCRs for gene therapy. Affinity-based methods include the use of barcoded peptide-MHC multimer libraries for profiling TCRs displayed on primary T cells^16,37,38^ and the assessment of soluble TCR interaction with peptide-MHC libraries displayed on the surface of yeast^18,39–41^. A potential limitation of such methods, however, is that they are unable to detect very low-affinity interactions between TCR and peptide-MHC that are nevertheless functional^42^. Epitope screening platforms based on cellular function include those relying on the display of peptide libraries by cells expressing chimeric MHC receptors^43,44^, cells undergoing trogocytosis^45^, or cells harboring reporters of exogenous granzyme activity^46,47^. Another method for the functional assessment of TCR cross-reactivity involves measuring T cell reactivity to single amino acid mutants of the target peptide, which has been recently utilized to profile the cross-reactivity of phage display-engineered TCRs reformatted for expression in primary T cells^48,49^ or a murine cell line^31^. While all of the above methods focus on specificity screening of TCRs, they have not been reported to directly enable TCR engineering.

Here we report the development and application of the <u>T</u>CR-accepti<u>n</u>g <u>T</u> cell (TnT) platform for the functional display, engineering and cross-reactivity profiling of TCRs. Reconstitution of Jurkat-derived TnT cells with transgenic TCRs was targeted by CRISPR-Cas9 to the endogenous TCRβ genomic region, specifically to the recombined complementary determining region 3 β (CDR3β) sequence, thus providing a monoallelic target that ensured a single integration event in every cell and physiological expression of transgenic TCRs. We use this approach to characterize > 30 individual TCRs and perform functional screening of 437 single amino acid variants and ~150,000 combinatorial variants. We dissect the mutational landscapes of two TCRs to reveal contrasting patterns and substantial discordance between antigen binding and antigen-induced signalling. Furthermore, through a combination of positive and negative functional selection steps coupled with deep sequencing, we identify combinatorial TCR variants with enhanced recognition of the MAGE-A3 cancer-testis antigen and lacking cross-reactivity to a known off-target peptide. Finally, we use peptide scanning to evaluate the cross-reactivity of resulting engineered TCRs displayed on TnT cells, leading to the identification of TCR variants targeting MAGE-A3 antigen with exquisite specificity and minimal cross-reactivity, which thus represent therapeutic candidates for TCR gene therapy.

## RESULTS

### Development of a <u>T</u>CR-accepti<u>n</u>g <u>T</u> cell (TnT) platform for the functional display of transgenic TCRs

The development of the TnT platform required extensive, multistep CRISPR-Cas9 genome editing. Using the human Jurkat E6-1 leukemia T cell line as a starting point, several genomic components were sequentially edited in order to facilitate the introduction and functional screening of TCRs of different specificities (Fig. 1a and Supplementary Fig. 1). Each genome editing step consisted of transfection with a gene-targeting guide RNA (gRNA) and, when required, a homology-directed repair (HDR) template encoding desired transgenes. This was followed by single-cell fluorescence-activated cell sorting (FACS), cell expansion and clone validation by flow cytometry and Sanger sequencing. In the first step, we equipped the Jurkat T cell line with constitutive Cas9 and human CD8 expression by CRISPR-Cas9 HDR targeting the CCR5 safe harbor locus^50^ (Supplementary Figs. 2a-e). This was performed in order to simplify and increase genome editing efficiency^51^, and to allow screening of CD8+ T cell-derived TCRs recognizing MHC class I-restricted peptides. In line with this, in the second step we knocked out the endogenous Jurkat CD4 co-receptor by CRISPR-Cas9 non-homologous end joining (NHEJ) (Supplementary Figs. 3a-c). In the third step, we introduced an NFAT-GFP construct into the AAVS1 safe harbor locus through CRISPR-Cas9 HDR, which provides a fluorescence reporter of TCR signaling and activation (Supplementary Figs. 4a-e). Notably, our design incorporated a promoter-less mRuby cassette that acted as a PPP1R12C gene-trap^52^ and served to identify successfully edited cells. In the fourth step, we targeted the endogenous Jurkat TCRα chain for knockout through CRISPR-Cas9 NHEJ, leading to the generation of a cell line with abolished surface expression of the TCR-CD3 complex (Supplementary Figs. 5a-f). In addition to eliminating the possibility of transgenic TCR chains mispairing with the endogenous Jurkat TCRα chain, this approach allows us to use restoration of CD3 surface expression as a selectable marker for successful integration of transgenic TCRs (Figs. 1a-c). In the final step, we knocked out expression of the Fas cell surface death receptor (Fas) by CRISPR-Cas9 NHEJ in order to provide our platform with resistance to activation-induced cell death (AICD) (Supplementary Figs. 6a-c). This resulting cell line constitutively expresses Cas9, human CD8 and mRuby, harbors an NFAT-GFP reporter of TCR signaling, and lacks expression of CD4, endogenous TCR and Fas, and thus represents the TnT platform used throughout the rest of this study.

**Figure 1.**
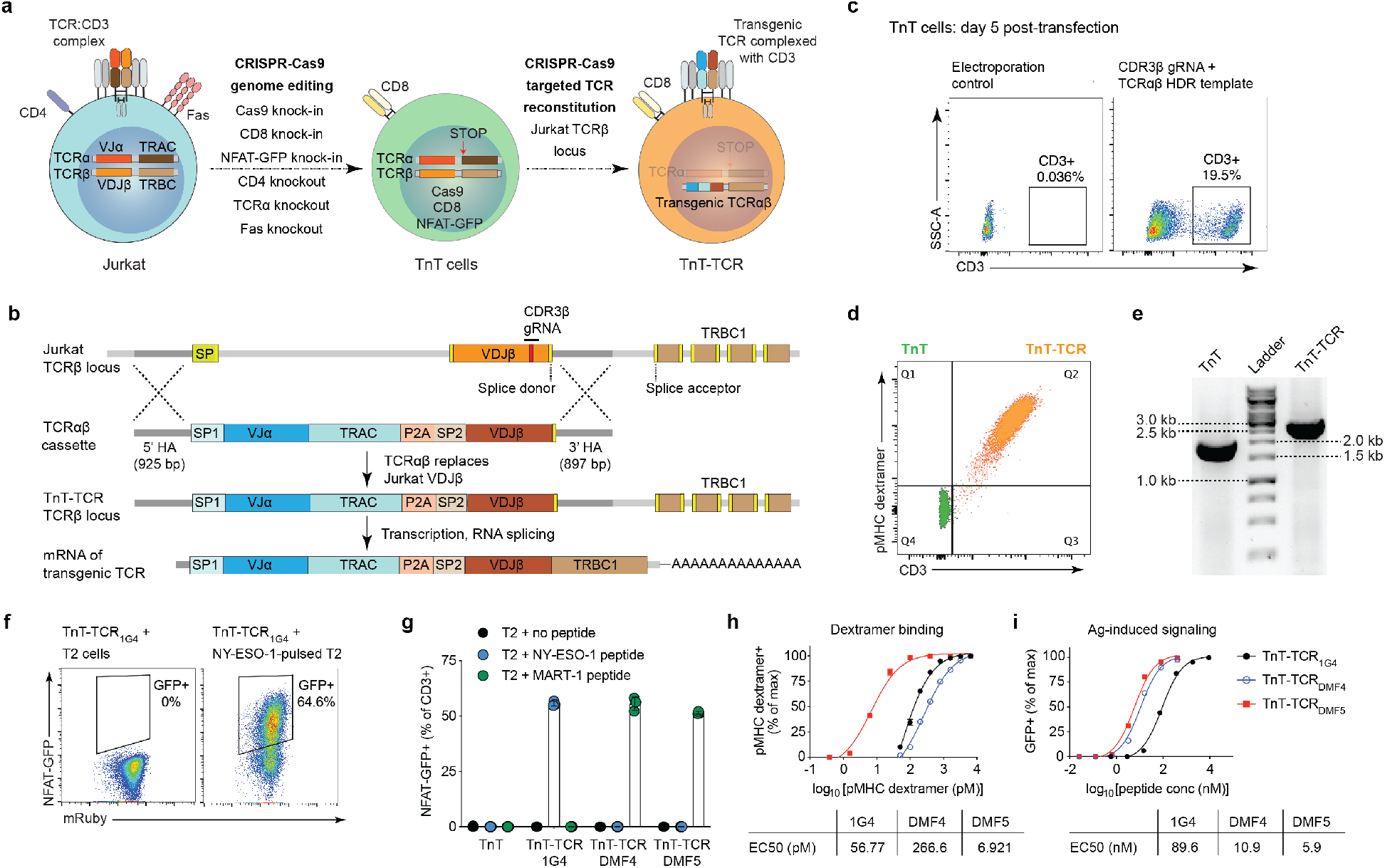
The TCR-accepting T cell (TnT) platform supports targeted reconstitution and functional display of transgenic TCRs. **a**, The Jurkat E6-1 cell line was subjected to sequential CRISPR-Cas9 genome editing steps in order to generate the TnT platform. TnT cells constitutively express Cas9 and human CD8, harbor an NFAT-GFP reporter of TCR signaling, and lack expression of CD4, Fas and endogenous TCR. Reconstitution of TnT cells with transgenic TCRs via CRISPR-Cas9 HDR results in TnT-TCR cells with restored surface expression of the TCR:CD3 complex. **b**, TCR reconstitution of TnT cells targeted to the Jurkat TCRβ locus. An HDR template encoding transgenic TCRαβ chains is integrated into TnT cells via Cas9 and a gRNA targeting the endogenous Jurkat CDR3β sequence. Transgenic TCR expression is dependent on correct RNA splicing with endogenous Jurkat TRBC1. **c**, Flow cytometric assessment of CD3 restoration in TnT cells after their targeted reconstitution with TCR_1G4_ (specific for NY-ESO-_157-165_ peptide). **d**, Representative flow cytometry plot showing peptide-MHC dextramer binding to TnT cells (no TCR expression) and MART-1-specific TnT-TCR_DMF5_ cells. **e**, Validation of targeted TCR reconstitution in TnT-TCR_DMF5_ cells by genomic PCR of the Jurkat VDJβ region. **f**, Representative flow cytometry dot plots displaying NFAT-driven GFP expression in TnT-TCR_1G4_ cells, but not TnT cells, after overnight co-culture with T2 cells pulsed with NY-ESO-1_157-164_ peptide. **g**, NFAT-GFP expression in TnT and TnT-TCR cells after overnight co-culture with T2 cells pulsed with NY-ESO-1_157-164_, MART-1_26-35(27L)_ or no peptide. **h**, Serially-diluted target peptide-MHC dextramers were used to assess the binding avidities of TnT-TCR_1G4_, TnT-TCR_DMF4_ and TnT-TCR_DMF5_ (n = 3). Peptide-MHC dextramer concentrations resulting in half-maximal proportions of dextramer positive cells (EC50) were derived from non-linear least squares fits. **i**, Normalized NFAT-GFP expression in TnT-TCR cells after overnight co-culture with T2 cells pulsed with serially-diluted cognate peptide (n = 2). Peptide pulse concentrations resulting in half-maximal proportions of NFAT-GFP+ cells (EC50) were derived from non-linear least squares fits. Data are displayed as mean ± SD.

We next reconstituted TnT cells with transgenic TCRs in a manner that would allow for monoallelic, homogenous and physiological TCR expression. To this end, we used CRISPR-Cas9 HDR to target integration of TCR transgenes to the recombined Jurkat TCRβ locus. First, we confirmed that the Jurkat T cell line expresses a single TCRβ chain by performing template-switching RT-PCR and Sanger sequencing (Supplementary Figs. 7a-b). We then designed and validated a gRNA molecule targeting the CDR3β of the Jurkat TCRβ chain (Supplementary Fig. 7c). Since the CDR3β sequence arises from the allele-independent recombination of V-, D- and J-genes, it provides a genomic target that is both highly specific and monoallelic. Having identified a suitable gRNA, we then proceeded to design TCRαβ HDR templates for targeted TCR reconstitution in TnT cells. We selected three previously discovered TCRs recognizing HLA-A*0201-restricted tumor-associated antigens: TCR_1G4_ with specificity to NY-ESO-1, and TCR_DMF4_ and TCR_DMF5_ with specificity to MART-1^53^ (Supplementary Table 1). HDR templates contained sequences encoding TCRα variable (VJα) and constant (TRAC) domains, a self-processing T2A peptide and a TCRβ variable (VDJβ) domain, flanked by ~ 900 bp homology arms mapping to the recombined Jurkat TCRβ locus (Fig. 1b). The lack of a constant TCRβ domain (TRBC) in the designed constructs made splicing with endogenous Jurkat TRBC exons a requirement for transgenic TCR expression. This feature allowed us to detect targeted genomic integration based on restored surface expression of CD3 following transfection of TnT cells with CDR3β gRNA and designed HDR templates (PCR product) (Fig. 1c). Furthermore, our strategy ensured that cells displaying restored CD3 expression underwent knockout of the endogenous Jurkat TCRβ chain, as integration of transgenic TCRs relied on the introduction of a dsDNA break at the targeted CDR3β genomic region. Although our initial experiments yielded HDR efficiencies of 1-2%, the development of an enhanced transfection protocol led to HDR efficiencies in the range of 5-20% (Fig. 3c and Supplementary Fig. 8). Targeted TCR reconstitution for the generation of TnT-TCR cells was further validated by detecting binding to cognate peptide-MHC dextramer using flow cytometry (Fig. 1d), PCR amplification of the Jurkat TCR genomic locus (Fig. 1e) and RT-PCR using reverse primers annealing to endogenous Jurkat TRBC sequences (Supplementary Figs. 9a and 9b). We thus successfully generated TnT-TCR_1G4_, TnT-TCR_DMF4_ and TnT-TCR_DMF5_ cell lines that were then subjected to further functional validation.

In order to assess antigen-induced signaling in TnT-TCR cells, we performed co-culture experiments using the HLA-A*0201-positive T2 cell line^54^. Co-culture of TnT-TCR_1G4_ cells with unpulsed (no peptide) T2 cells yielded no detectable expression of NFAT-GFP, while co-culture with T2 cells pulsed with NY-ESO-1_157-165_ cognate peptide induced robust expression of the NFAT-GFP reporter (Fig. 1f). To further assess the specificity of our platform, we performed co-cultures of TnT, TnT-TCR_1G4_, TnT-TCR_DMF4_ and TnT-TCR_DMF5_ cells with T2 cells pulsed with NY-ESO-1_157-165_ peptide, MART-1_26-35(2L)_ peptide or no peptide. We found that NFAT-GFP expression was fully restricted to correct TCR-peptide pairings, with no detectable NFAT-GFP expression across negative controls (Fig. 1g).

We next compared the binding and functional avidities of TnT-TCR_1G4_, TnT-TCR_DMF4_ and TnT-TCR_DMF5_ cells (Fig. 1h-i). TnT-TCR_DMF5_ cells displayed the highest binding avidity to their target peptide-MHC (EC50 = 7 pM), followed by TnT-TCR_1G4_ (EC50 = 57 pM) and TnT-TCR_DMF4_ (EC50 = 267 pM), with picomolar EC50 values reflecting the multivalent nature of surface-displayed TCRs and peptide-MHC dextramers (10-20 peptide-MHC copies per molecule). We evaluated the functional avidity of TnT-TCR cells following co-culture with T2 cells pulsed with serial dilutions of cognate peptide, which revealed a dose-dependent response in terms of NFAT-GFP-expressing cells (Fig. 1i). TnT-TCR_DMF5_ displayed the highest functional avidity (EC50 = 6 nM), followed by TnT-TCR_DMF4_ (EC50 = 11 nM) and TnT-TCR_1G4_ (EC50 = 90 nM). In contrast to their 39-fold difference in binding to peptide-MHC dextramer, the difference between TnT-TCR_DMF4_ and TnT-TCR_DMF5_ in terms of functional avidity was only 2-fold. This indicated that a high binding avidity is not a requirement for antigen-induced expression of NFAT-GFP in TnT-TCR cells. Finally, additional co-culture experiments confirmed TnT-TCR resistance to AICD (Supplementary Fig. 6d) and physiological down-regulation of surface TCR-CD3 expression with increasing amounts of presented antigen^56^ (Supplementary Fig. 6e).

### Deep mutational scanning reveals the expression, binding and functional landscapes of TCRs

In contrast to previous TCR engineering methods^28,29^, the TnT platform allows us to assess TCRs across multiple parameters which include their surface expression in complex with CD3, binding to peptide-MHC multimers and signaling in response to antigen-presentation. We selected the MART-1-specific TCR_DMF4_ and the MAGE-A3-specific TCR_A3_ for comprehensive profiling using the TnT platform (Fig. 2a). TCR_A3_ is a low-avidity TCR isolated from a melanoma patient treated with a viral vaccine encoding the HLA-A*0101-restricted MAGE-A3_168-176_ peptide^57,58^ (Supplementary Table 1). Co-culture experiments with the MAGE-A3-positive EJM myeloma cell line revealed that the avidity of TCR_A3_ was not sufficient to induce NFAT-GFP expression in TnT-TCR_A3_ cells. However, we detected increased surface expression of the early T cell activation marker CD69 following co-culture, which provided us with a complementary and more sensitive functional readout (Supplementary Fig. 10c).

**Figure 2.**
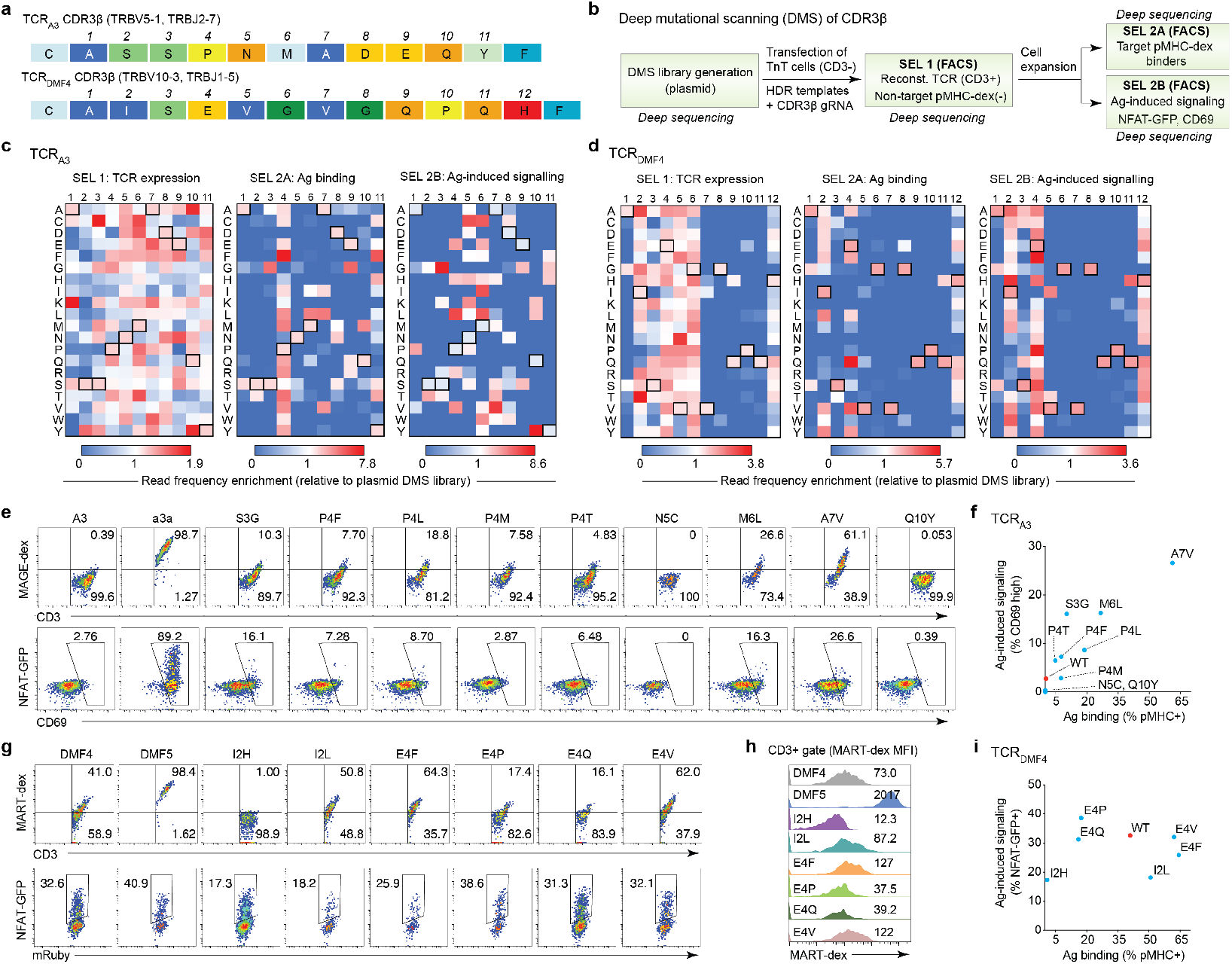
Deep mutational scanning of CDR3β reveals the sequence landscape for TCR expression, binding and signaling. **a**, The CDR3β sequences of TCR_A3_ and TCR_DMF4_ were subjected to deep mutational scanning (DMS). **b**, DMS libraries were generated by means of plasmid nicking saturation mutagenesis and integrated into TnT cells by Cas9 HDR. Reconstituted cells were selected by FACS based on CD3 surface expression, target peptide-MHC binding and antigen-induced signalling (co-culture with antigen-presenting cells). Deep sequencing of CDR3β sequences was performed at every selection step. **c**, **d**, Heatmaps displaying the enrichment of sequencing reads from TCR_A3_ and TCR_DMF4_ variants across selections relative to their occurrence in the starting plasmid DMS libraries. Outlined boxes represent wild-type CDR3β residues. **e-f**, Flow cytometric assessment of antigen binding and antigen-induced signalling in TnT cells reconstituted with TCR_A3_, TCR_a3a_ and selected TCR_A3_ variants (CD3+ gate). **e**, The levels of binding to MAGE-A3 peptide-MHC dextramer (top row) and activation after overnight co-culture with MAGE-A3+ EJM cells (bottom row, proportion CD69^high^ TnT-TCR) are displayed. **f**, Graph displays the levels of antigen binding and antigen-induced signaling in TCR_A3_ single mutants relative to wild-type TCR_A3_. **g-i**, Flow cytometric assessment of antigen binding and antigen-induced signalling in TnT cells reconstituted with TCR_DMF4_, TCR_DMF5_ and selected TCR_DMF4_ variants (CD3+ gate). **g**, The levels of binding to MART-1 peptide-MHC dextramer (top row) and activation after overnight co-culture with MART-1-pulsed T2 cells (bottom row, proportion NFAT-GFP+ TnT-TCR) are displayed. **i**, Histograms displaying the levels of MART-1 peptide-MHC dextramer bound by TnT-TCR cells. **h**, Graph displays the levels of antigen binding and antigen-induced signaling in TCR_DMF4_ single mutants relative to wild-type TCR_DMF4_.

Having characterized parental TCR_DMF4_ (Fig. 1g-i) and TCR_A3_, we aimed to determine their mutational landscapes across the multiple parameters provided by TnT cells. To this end, we performed deep mutational scanning (DMS) of CDR3β as this TCR region is typically enriched for direct contacts to peptide antigen rather than to MHC^55^. To generate DMS libraries we introduced NNK degenerate codons at each CDR3β position using plasmid nicking saturation mutagenesis^59^ (Supplementary Figs. 11a-c). Plasmid DMS libraries were then used to generate TCRαβ HDR templates (PCR products), which were transfected into TnT cells alongside CDR3β gRNA (Supplementary Fig. 11b). TnT cells displaying restored TCR-CD3 surface expression were isolated by FACS (SEL 1), expanded and re-sorted based on binding to cognate peptide-MHC dextramer (SEL 2A) or activation following co-culture with cells displaying target peptides (SEL 2B) (Fig. 2b and Supplementary Figs. 12a-c). Due to the low avidity of the parental TCR_A3_, functional selections of TCR_A3_ DMS libraries were based on CD69 expression, instead of NFAT-GFP. Following FACS, deep sequencing was performed to identify TCR variants that were enriched across selection steps.

Sequence enrichment analysis was performed relative to observed variant frequencies in plasmid DMS libraries of TCR_A3_ (Fig. 2c) and TCR_DMF4_ (Fig. 2d). We found that wild-type TCR sequences (Figs. 2c and 2d, boxed residues) were enriched in most selections, which likely reflected the depletion of unfavorable variants. The only exception occurred in TCR_A3_ SEL 2B (signaling), in which wild-type TCR_A3_ was depleted relative to the original plasmid library. This was in agreement with the low levels of antigen-induced signaling found in TnT-TCR_A3_ cells (Supplementary Fig. 12c), and pointed towards the occurrence of TCR_A3_ single mutants with substantial increases in function relative to wild-type TCR_A3_. In terms of TCR-CD3 expression (SEL 1), we detected sequencing reads originating from all but five TCR_A3_ variants, indicating that the vast majority of TCR_A3_ mutants were capable of cell surface expression in complex with CD3 (Fig. 2c). This was in stark contrast to the TCR_DMF4_ surface expression landscape, which was heavily restricted (Fig. 2d). As expected, the mutational landscapes for peptide-MHC binding (SEL 2A) and antigen-induced signaling (SEL 2B) were more restricted than those seen for TCR-CD3 surface expression (Figs. 2c and 2d). In the case of TCR_A3_, while a similar number of enriched variants was observed in binding (SEL 2A) and signaling (SEL 2B) selections, there was little correlation between the deep sequencing enrichment levels of individual variants in the two fractions (Fig. 2c and Supplementary Fig. 13a). Some TCR_A3_ variants, however, did display high levels of sequence enrichment for both parameters (e.g., S3G, P4L, M6L and A7V) (Supplementary Fig. 13a). In the case of TCR_DMF4_, there was a modest correlation between enrichment levels of individual variants in binding (SEL 2A) and signaling (SEL 2B) fractions (Supplementary Fig. 13b). However, a very limited number of TCR_DMF4_ variants displayed enrichment levels that were higher than wild-type TCR_DMF4_ for either binding or signaling. Notably, the number of enriched TCR_DMF4_ variants in the signaling fraction (SEL 2B) was considerably higher than in the binding fraction (SEL 2A). This indicated that antigen-induced signaling may occur in the absence of detectable peptide-MHC binding in some of these TCR_DMF4_ variants, particularly those with substitutions in CDR3β positions 2-4 (Fig. 2d).

In order to further characterize the discordance observed between antigen binding and antigen-induced signaling, we reconstituted TnT cells with variants showing enrichment above wild-type TCR levels. For TCR_A3_, we selected nine variants (S3G, P4F, P4L, P4M, P4T, N5C, M6L, A7V, Q10Y) and included the high-affinity TCR_a3a_ as a positive control (see next section for more information on TCR_a3a_). For TCR_DMF4_, we selected six variants (I2H, I2L, E4F, E4Q, E4P, E4V) and included the high-affinity TCR_DMF5_ as a positive control. All selected TCR variants were successfully expressed on the surface of TnT cells and were assessed for binding with cognate peptide-MHC dextramer (Fig. 2e, g top row) and antigen-induced signaling (Fig. 2e, g bottom row). Notably, the majority of TCR_A3_ variants displayed enhancements in binding and signaling relative to wild-type TCR_A3_. Consistent with sequence enrichment data, variants S3G, P4L, M6L and A7V displayed the largest increases in terms of both binding and signaling (Fig. 2f). For TCR_DMF4_ variants, only modest improvements were observed relative to wild-type (Fig. 2i), namely enhanced binding of variants I2L, E4V and E4F to MART-1 peptide-MHC dextramer (Fig. 2 h). Surprisingly, TCR_DMF4_ variants I2H and I2L showed similar levels of antigen-induced signaling despite I2H showing no detectable binding to MART-1 peptide-MHC dextramer (Figs. 2g and 2h), further highlighting the discordance between binding and signaling in certain TCRs.

### Functional selection of combinatorial CDR3β libraries identifies TCRs with enhanced target specificity and reactivity

Next, we aimed to engineer the low-avidity TCR_A3_ for enhanced reactivity to the cancer-testis antigen MAGE-A3, as it represents an attractive target that is widely expressed in several types of epithelial tumors^60^. In a previous effort, TCR_A3_ was engineered by phage display to generate TCR_a3a_, a variant with >200-fold higher affinity to MAGE-A3_168-176_ peptide-MHC^14,30^ (Supplementary Fig. 10a and Supplementary Table 1). However, use of TCR_a3a_ in a gene therapy clinical trial resulted in two treatment-induced patient deaths^14^. Retrospective analysis uncovered that TCR_a3a_ cross-reactivity to a peptide derived from the protein titin, which is highly expressed in beating cardiomyocytes, led to fatal cardiac toxicity^30^. Notably, this occurred despite the titin-derived peptide (ESDPIVAQY) having four amino acid differences relative to MAGE-A3_168-176_ (EVDPIGHLY). Consistent with previous findings, we found that TnT-TCR_A3_ cells showed low but detectable binding and TnT-TCR_a3a_ showed high binding to MAGE-A3 peptide-MHC dextramer, while only TnT-TCR_a3a_ displayed binding to titin peptide-MHC dextramer (Supplementary Fig. 10b).

We sought to generate and screen combinatorial CDR3β libraries using the TnT platform in order to identify TCR variants with enhanced antigen specificity and reactivity. Combinatorial library design was based on data obtained from TCR_A3_ DMS (Fig. 2c), as this allowed us to maximize both library functionality and size for screening in mammalian cells^51^. By utilizing a sequence space optimization algorithm^51^, we designed degenerate codons that mimic the TCR_A3_ single mutant frequencies found in DMS binding and signaling selections (Supplementary Fig. 14), leading to a combinatorial library diversity of 2.6 x 10^5^ (Fig. 3a). The designed TCR_A3_ combinatorial library was generated by overlap-extension PCR, and cloned into a TCR_A3_-encoding plasmid for bacterial transformation (Supplementary Fig. 15a). We estimated that the physical plasmid library contained as many as 1.5 x 10^5^ TCR_A3_ variants based on the number of bacterial transformants and the proportion of unique clones after deep sequencing (Supplementary Fig. 15b). The constructed TCR_A3_ plasmid library was used to generate HDR templates, which were then integrated into TnT cells by CRISPR-Cas9 HDR.

**Figure 3.**
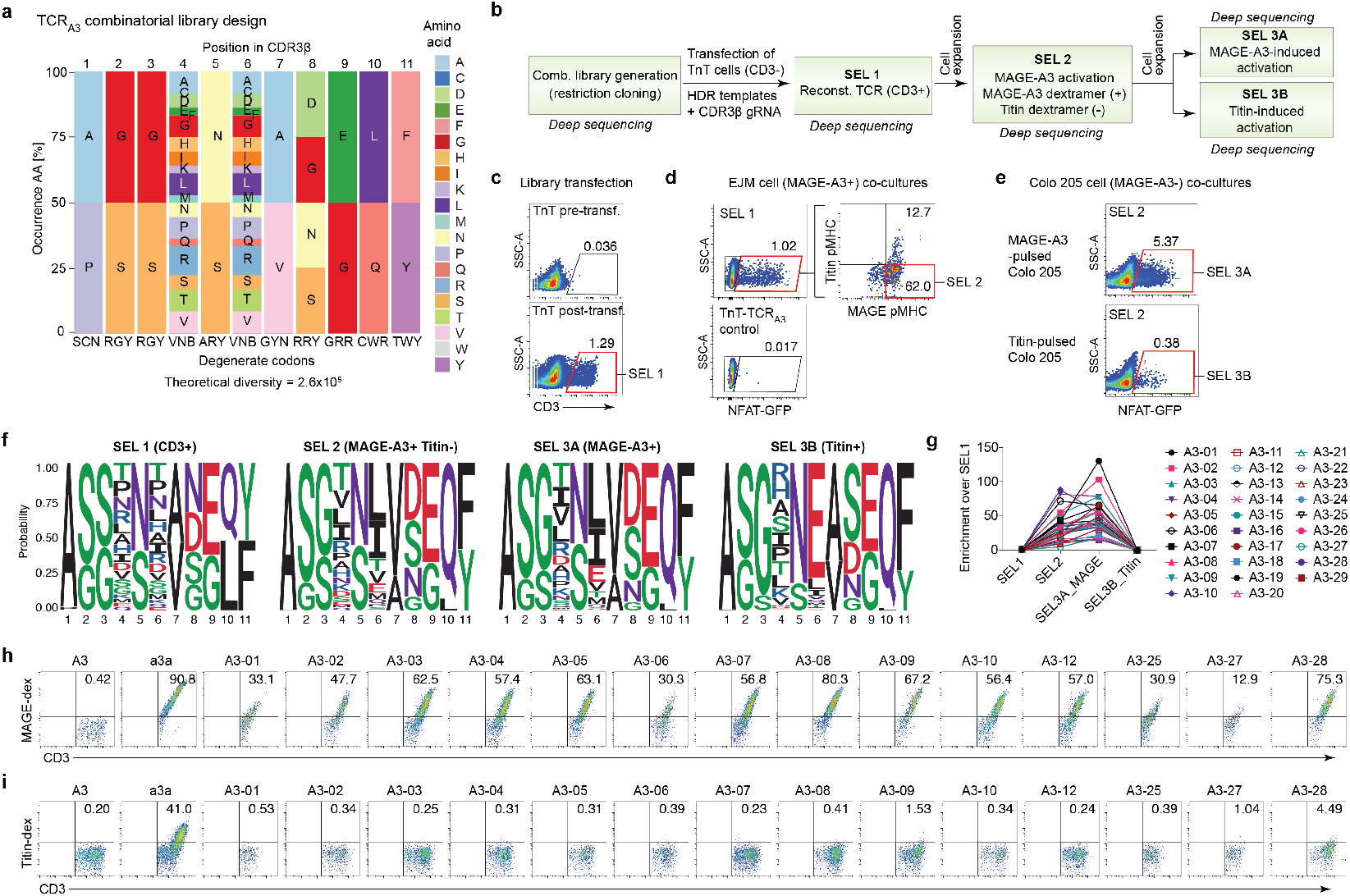
Rational design and functional screening of combinatorial TCR mutagenesis libraries in TnT cells for selection of enhanced target specificity. **a**, Data obtained from TCR_A3_ CDR3β DMS was utilized as input to design a combinatorial mutagenesis library that has a theoretical diversity of 2.6 x 10^5^ variants. Degenerate codons were designed to recapitulate the amino acid frequencies observed in DMS selections based on MAGE-A3 peptide-MHC dextramer binding and MAGE-A3-induced signalling. **b**, Strategy for the selection of TCR_A3_ combinatorial mutagenesis variants with enhanced recognition of the MAGE-A3_168-176_ peptide (EVDPIGHLY), while avoiding cross-reactivity to the known titin_24,337-24,345_ off-target peptide (ESDPIVAQY). Deep sequencing of CDR3β sequences was performed at every selection step. **c**, Flow cytometry plots show selection of TCR_A3_ variants from the combinatorial library that are capable of surface expression in TnT cells. TnT cells with restored CD3 surface expression were bulk-sorted (SEL 1). **d**, Flow cytometry plots showing TnT-TCR cells (from SEL 1) with positive MAGE-A3-induced signaling (NFAT-GFP+), positive MAGE-A3 peptide-MHC binding and negative titin peptide-MHC binding; cells were bulk-sorted (SEL 2) and expanded in culture. **e**, Expanded SEL 2 cells were co-cultured overnight with either MAGE-A3 peptide-pulsed or titin peptide-pulsed Colo 205 cells and bulk-sorted for activation by NFAT-GFP+ expression (SEL 3A and SEL 3B). **f**, Amino acid sequence logos showing the frequency of specific residues at each CDR3β position across selections (logos weighted on unique clone frequencies). **g**, TCR_A3_ variants with predicted high specificity for MAGE-A3 and their enrichment across selections based on deep sequencing data. **h-i**, Flow cytometry plots of TnT cells reconstituted with TCR_A3_, TCR_a3a_ and selected TCR_A3_ combinatorial variants. Cells were assessed for CD3 expression, MAGE-A3 peptide-MHC binding (**h**) and titin peptide-MHC binding (**i**). Cells in the CD3+ gate are shown. Degenerate nucleotide symbols: R = A, G; Y = C, T; S = G, C; W = A, T; K = G, T; M = A, C; B = C, G, T; D = A, G, T; H = A, C, T; V = A, C, G; N = any base.

The selection strategy for the TnT-TCR_A3_ library consisted of a combination of positive and negative selection steps based on binding to peptide-MHC dextramers and signaling in response to cell-displayed antigens, with deep sequencing of TCR regions performed after every selection step (Fig. 3b). In the first selection step, TnT cells displaying restored CD3 expression were isolated by FACS and expanded (SEL 1, Fig. 3c). Next, expanded cells from SEL 1 were co-cultured overnight with MAGE-A3-positive EJM cells and co-stained with MAGE-A3 and titin peptide-MHC dextramers. Different to wild-type TnT-TCR_A3_ cells, a fraction of TnT-TCR_A3_ library cells displayed robust NFAT-GFP expression (Fig. 3d). Analysis of the NFAT-GFP-positive population revealed that ~ 20% of cells that bound to MAGE-A3 also recognized the titin peptide-MHC dextramer (Fig. 3d). In order to exclude these cross-reactive variants, we isolated cells that were NFAT-GFP-positive, MAGE-A3 peptide-MHC dextramer-positive and titin peptide-MHC dextramer-negative (SEL 2). A final functional selection step was performed by co-culturing expanded cells from SEL 2 with Colo 205 cells pulsed with either MAGE-A3 or titin peptides. As expected, we observed a substantial fraction of TnT-TCR cells with NFAT-GFP expression after co-culture with MAGE-A3-pulsed Colo 205 cells (Fig. 3e). Interestingly, we also found that a subset of TnT-TCR cells still expressed NFAT-GFP in response to titin stimulation despite the exclusion of titin peptide-MHC binders in SEL 2. Analysis of the sequence space landscape revealed a number of notable trends, including the preference of alanine codons at position CDR3β-1 for TCR:CD3 expression (SEL 1), and the increased frequency of specific residues at CDR3β-6 and CDR3β-10 in MAGE-A3-based selections (SEL 2 and SEL 3A). Most notably, we observed a substantial increase in the frequency of glutamate residues at CDR3β-6 in SEL 3B (titin-induced signaling), highlighting this substitution as a potential determinant of titin cross-reactivity.

We next analyzed deep sequencing data for clone enrichment (Supplementary Table 2). We identified a total of 195 unique clones displaying greater than 2-fold enrichment in SEL 3A (MAGE-A3 signaling) and that were either absent or not enriched in SEL 3B (titin signaling). We scored these clones based on their frequency and enrichment levels and identified the top 29 candidate TCR_A3_ variants, of which we selected 14 for further characterization (Fig. 3g and Supplementary Table 3). To this end, we reconstituted TnT cells with wild-type TCR_A3_, TCR_a3a_ and selected TCR_A3_ variants via CRISPR-Cas9 HDR and assessed them for CD3 expression, binding to MAGE-A3 and titin peptide-MHC dextramers. All selected TCR_A3_ variants were successfully displayed on the surface of TnT cells and showed enhanced MAGE-A3 dextramer binding relative to wild-type TnT-TCR_A3_ cells (Fig 3h). Importantly, binding to titin peptide-MHC dextramer was undetectable in most of the selected TCR_A3_ variants, while readily detected in TnT cells displaying the phage display-engineered TCR_a3a_ (Fig 3i).

### The TnT platform enables TCR cross-reactivity profiling and accurate prediction of off-target antigens

Since engineering TCRs for enhanced reactivity carries the risk of introducing unwanted specificities, we decided to apply the TnT platform for the screening of TCR cross-reactivity. As a proof of concept, we profiled the cross-reactivity of the phage display-engineered TCR_a3a_^30^, which has titin as a known off-target, by performing deep mutational scanning (DMS) of the MAGE-A3 target peptide. Accordingly, we designed a synthetic peptide DMS library containing every possible single amino acid mutant of MAGE-A3_168-176_ (EVDPIGHLY) (n = 171). Each library peptide was then individually pulsed on Colo 205 cells (HLA-A*0101-positive, MAGE-A3-negative) and co-cultured with TnT-TCR_a3a_ cells (n = 171 co-cultures). Activation of TnT-TCR_a3a_ cells was assessed by NFAT-GFP expression, which was corrected for background and normalized to the response induced by Colo 205 cells pulsed with wild-type MAGE-A3 peptide (Fig. 4a). In agreement with a previous glycine-serine scan^30^, we found that most mutations at peptide-MHC positions 1, 3, 4, 5 and 9 of the target MAGE-A3 peptide resulted in substantially reduced TnT-TCR_a3a_ activation. By contrast, several peptides with mutations at positions 2, 6, 7 and 8 induced strong activation of TnT-TCR_a3a_ cells.

**Figure 4.**
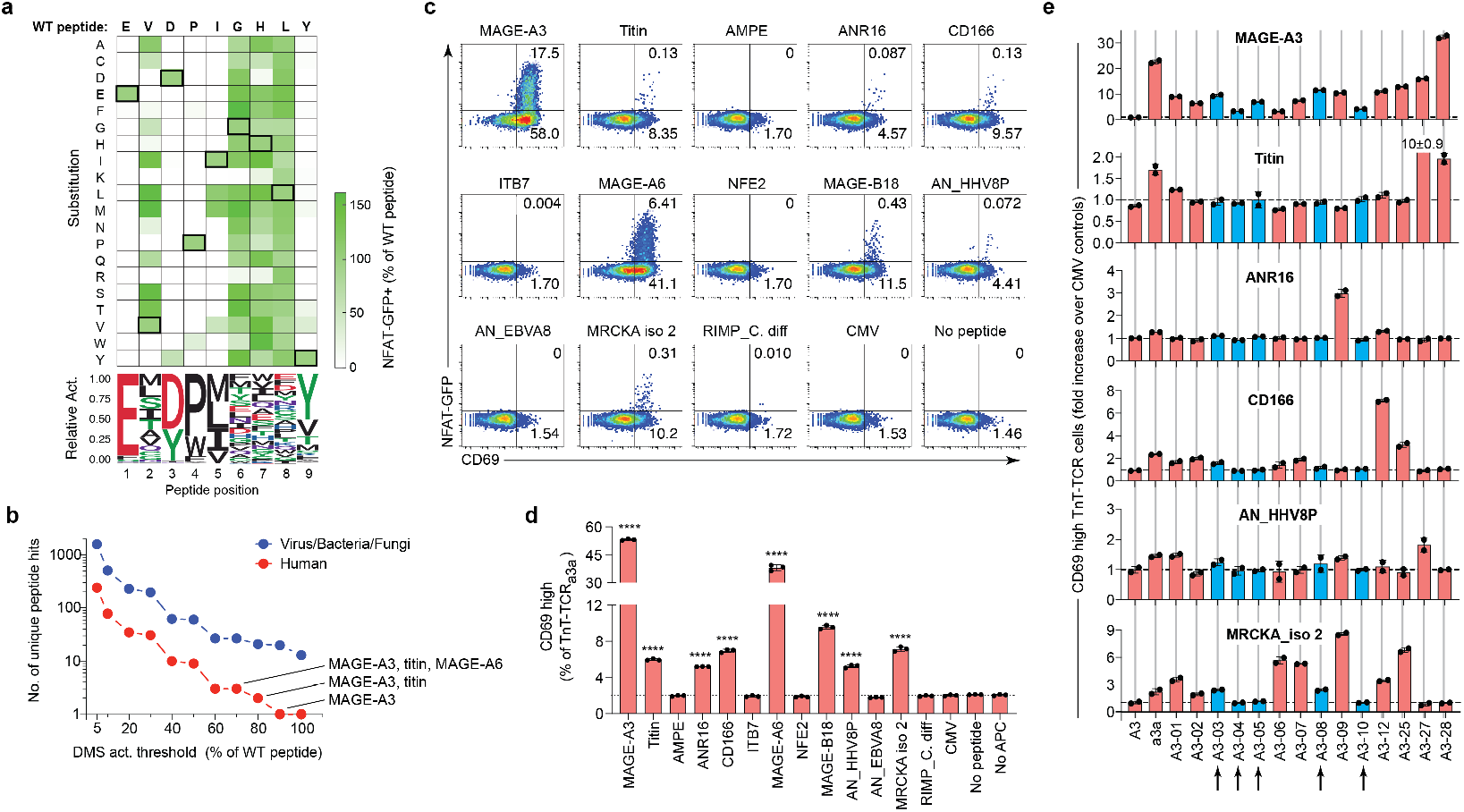
The TnT platform enables TCR cross-reactivity profiling and prediction of off-target peptides. **a**, The cross-reactivity profile of TnT-TCR_a3a_ cells was assessed using single mutant variants of the wild-type MAGE-A3 peptide (peptide DMS library), which were pulsed on Colo 205 cells for individual co-culture assays (n = 171). Heatmap shows the proportion of NFAT-GFP+ TnT-TCR_a3a_ cells after co-culture, as determined by flow cytometry. Data are normalized to the response induced by the MAGE-A3 wild-type peptide (boxed residues). Sequence logo shows the relative activity of peptide DMS library members carrying mutations at the same peptide position. **b**, The sequences of peptides mediating 5, 10, 20, 30, 40, 50, 60, 70, 80, 90 and 100 percent activation relative to the MAGE-A3 wild-type peptide were utilized to generate motifs to query the UniProtKB database. Dot plot displays the number of human (red) and non-human (blue) unique peptide hits resulting from these searches. The first (MAGE-A3), second (titin) and third (MAGE-A6) highest predicted activating peptides are highlighted. **c**, Flow cytometry of CD69 and NFAT-GFP expression following co-culture with peptide-pulsed Colo 205 cells shows cross-reactivity of TnT-TCR_a3a_ cells against a subset of predicted off-target peptides. The CMV peptide (HLA-A*0101-restricted, VTEHDTLLY) is included as a negative control. **d**, Bar graph shows repeat of the experiment in (**c**) performed in triplicate. The percentages of CD69^high^ TnT-TCR_a3a_ cells after co-culture with peptide-pulsed Colo 205 cells were utilized to assess reactivity to each peptide. Dotted line reflects the mean of CMV-pulsed controls (Y = 1.98). Asterisks indicate significant differences to CMV controls as determined by two-way ANOVA with Bonferroni post-hoc test for multiple comparisons. **e**, Assessment of selected TnT-TCR_A3_ CDR3β variants for cross-reactivity. TnT cells expressing TCR_A3_, TCR_a3a_ and selected TnTA3 variants were co-cultured with Colo 205 cells pulsed with a subset of predicted off-target peptides. The percentages of CD69^high^ TnT-TCR cells were determined by flow cytometry and normalized to their respective CMV backgrounds (n = 2). Selected TnT-TCR_A3_ variants displaying favorable cross-reactivity profiles are highlighted in blue. Data are displayed as mean ± SD. * *P* < 0.05, ** *P* < 0.01, *** *P* < 0.001, **** *P* < 0.0001, ns = not significant. See Supplementary Table 5 for a full list of peptides and their sequences.

In order to predict potential off-targets, we generated peptide sequence space motifs of allowed substitutions at discrete thresholds of TnT-TCR_a3a_ activation, and used them to interrogate the UniProtKB database^48,49^. As expected, the number of unique peptide hits resulting from these searches decreased with increasing activation thresholds (Fig. 4b), as this reflected the use of more restricted peptide motifs (Supplementary Table 4). The only human peptide returned at the highest activation threshold (100%) was indeed the target MAGE-A3_168-176_ (EVDPIGHLY) (Fig. 4b). Remarkably, the human peptide returned at the second highest activation threshold was the titin peptide ESDPIVAQY, which was identified despite the fact that the most similar peptides in the DMS library had three amino acid mismatches. The human peptide returned at the third highest activation threshold was MAGE-A6_168-176_ (EVDPIGHVY), another known target of TCR_a3a_^30^. We also identified the cancer-testis antigen MAGE-B18_166-174_ (EVDPIRHYY), which has been previously shown to activate TCR_a3a_^30^.

We next selected a subset of predicted off-target peptides with potential clinical relevance for experimental validation. These were peptides returned across several activation thresholds and included ten human and two viral peptides (Supplementary Table 5). We also included the bacterial *C. difficile* EKDPIKENY peptide, which was predicted in a previous study to be a potential off-target of TCR_a3a_^30^, but was not predicted in our high-resolution DMS analysis as inclusion of lysine at position 2 resulted in negligible TnT-TCR_a3a_ activation (Fig. 4a). Peptides were pulsed on Colo 205 cells and co-cultured with TnT-TCR_a3a_ cells overnight, followed by assessment of activation based on both NFAT-GFP and CD69 expression. MAGE-A3 induced strong expression of NFAT-GFP and CD69, while MAGE-A6 and MAGE-B18 also induced substantial but lower activation (Fig. 4c). Since NFAT-GFP expression was low in some co-cultures, quantification of CD69^high^ TnT-TCR_a3a_ cells allowed us to identify peptides inducing responses significantly higher than background (CMV negative control peptide) (Fig. 4d). We found that titin significantly activated TnT-TCR_a3a_ cells, however this response was still considerably lower than that induced by MAGE-A3, which is consistent with the reported 50-fold difference in the binding affinity of TCR_a3a_ towards titin peptide-MHC^30^. We also found that human peptides derived from the proteins ANR16, CD166 and MRCKA isoform 2; and a peptide derived from a Kaposi Sarcoma-associated herpesvirus exonuclease (AN_HHV8P) induced significant activation of TnT-TCR_a3a_ cells (Fig. 4d), and thus represent potentially novel and clinically relevant off-targets of TCR_a3a_.

Having profiled TCR_a3a_ cross-reactivity through target peptide DMS, we wondered if any of the resulting off-targets were capable of activating our engineered TCR_A3_ combinatorial variants (Fig. 3g-i). To assess this possibility, we co-cultured TnT cells expressing wild-type TCR_A3_, TCR_a3a_ or selected TCR_A3_ variants with Colo 205 cells pulsed with activating off-target peptides (excluding MAGE-A6 and MAGE-B18), as well as MAGE-A3, titin and CMV control peptides (Fig. 4e). We found that all TnT-TCR_A3_ variants displayed a stronger response to MAGE-A3 than wild-type TnT-TCR_A3_, which showed negligible activation. Interestingly, we found that TnT-TCR_A3-27_ and TnT-TCR_A3-28_ were activated by titin. While the low response observed in TnT-TCR_A3-28_ cells might be explained by its residual level of binding to titin peptide-MHC dextramer (Fig. 3i), the robust response in TnT-TCR_A3-27_ cells was highly unexpected considering its minimal binding to titin peptide-MHC dextramer (Fig. 3i). In contrast, TnT-TCR_A3-09_ showed higher binding to titin peptide-MHC dextramer than TnT-TCR_A3-27_ cells (Fig 3i) but were not activated in response to titin-induced signaling (Fig 5e and Supplementary Fig. 15). These results provide further examples of the discordance between TCR antigen binding and signaling, and emphasize the importance of functional screening to better assess TCR specificity and cross-reactivity. Another interesting finding was that negative selection against titin recognition (Figs. 3b and 3c) did not impede the emergence of additional cross-reactivity to other peptides in some of the engineered TCR_A3_ variants. This was clearly the case for variants TCR_A3-09_ (cross-reactive to ANR16 and MRCKA) and TCR_A3-12_ (cross-reactive to CD166 and MRCKA), which provide examples of increased cross-reactivity following TCR engineering (Fig. 4e). Most importantly, we found that variants TCR_A3-03_, TCR_A3-04_, TCR_A3-05_, TCR_A3-08_ and TCR_A3-10_ showed low or negligible activation in response to all tested peptides (Fig. 4e, highlighted in blue, and Supplementary Fig. 17).

**Figure 5.**
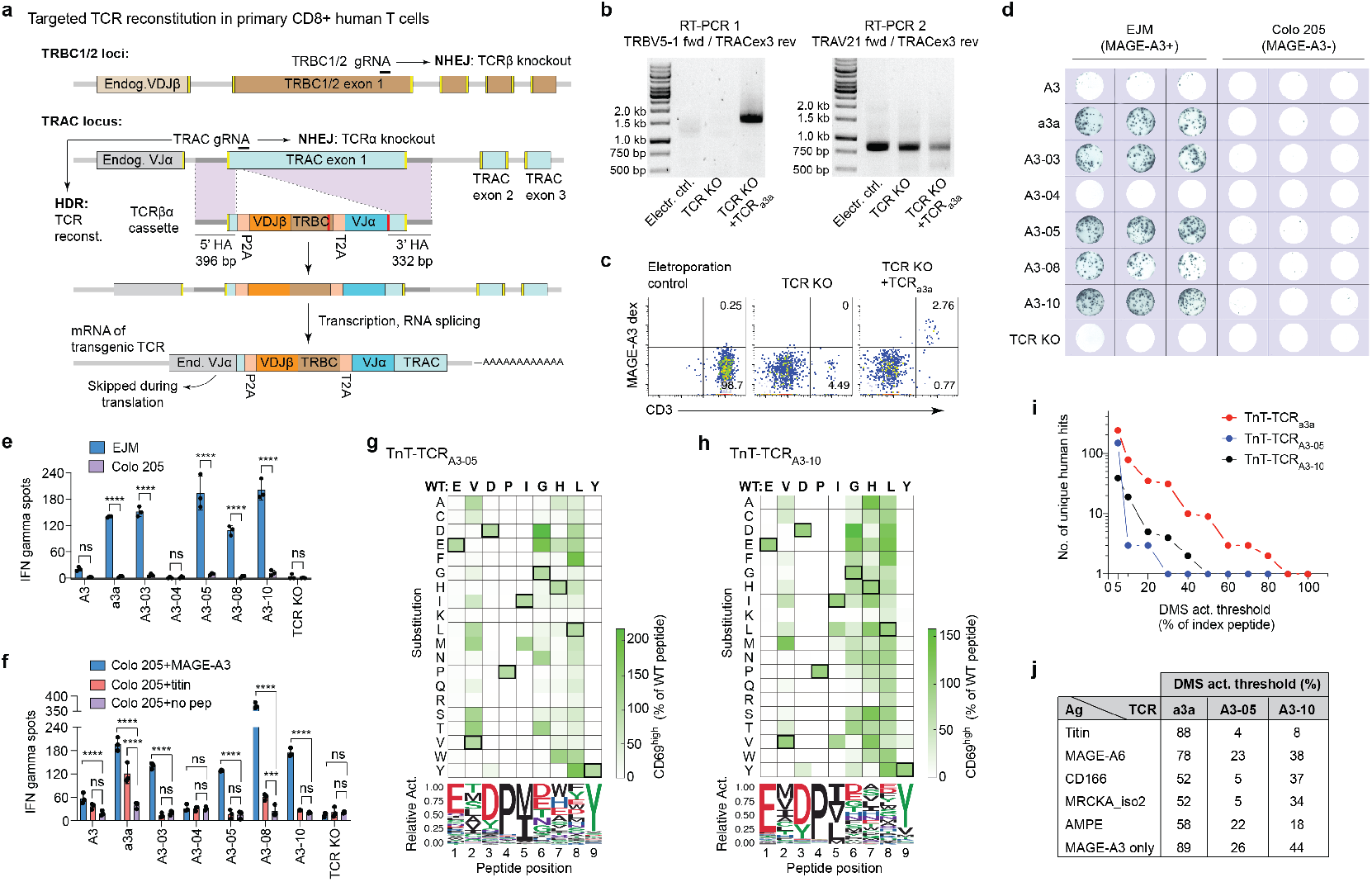
The TnT platform and primary T cells validate the engineering of TCRs with enhanced target reactivity and minimal cross-reactivity. **a**, Schematic shows TCR reconstitution in primary human CD8+ T cells is targeted to the TRAC locus by Cas9 HDR. T cells were transfected with TRAC and TRBC1/2 gRNAs in complex with recombinant Cas9 for dual knockout of endogenous TCRα and TCRβ chains. TCRβα cassettes consisted of a P2A self-processing peptide, followed by VDJβ and TRBC domains, a T2A self-processing peptide and a VJα domain. Regions highlighted in red were re-coded in order to prevent targeting of HDR templates. Transgenic TCR expression is dependent on correct RNA splicing with endogenous TRAC exons 2 and 3. **b**, Validation of targeted TCR_a3a_ integration by means of RT-PCR with a forward primer annealing to the transgenic TRBV5-1 gene and a reverse primer annealing to the endogenous TRAC exon 3. Electroporation control and dual TCR knockout samples display no PCR product, while samples derived from TCR_a3a_ transfectants display a 1.6 kb product consistent with targeted TCR integration and correct splicing with endogenous TRAC. A second RT-PCR reaction amplifying both endogenous and transgenic TRAV21-TRAC sequences was included as a positive control. **c**, Flow cytometric assessment of CD3 expression and MAGE-A3 peptide-MHC dextramer binding in primary CD8+ T cells reconstituted with TCR_a3a_. **d**, IFN-γ ELISPOT assays for assessment of primary CD8+ T cells reconstituted with selected TCRs following co-culture with MAGE-A3-positive EJM cells or MAGE-A3-negative Colo 205 cells (n = 3; 5×10^4^ T cells per well). **e**, Quantification of IFN-γ ELISpot data in (**d**). Asterisks indicate significant differences as determined by two-way ANOVA with Bonferroni post-hoc test for multiple comparisons. **f**, Quantification of IFN-γ ELISpot data from primary CD8+ T cells reconstituted with selected TCRs following overnight co-culture with Colo 205 cells pulsed with MAGE-A3_168-176_, titin_24,337-24,345_, or no peptide (n = 3; 4×10^5^ T cells per well). Asterisks indicate significant differences to non-pulsed Colo 205 controls as determined by two-way ANOVA with Bonferroni post hoc test for multiple comparisons. **g**, **h**, The cross-reactivity profiles of TnT-TCR_A3-05_ and TnT-TCR_A3-10_ cells were assessed by individual co-cultures with Colo 205 cells pulsed with single mutant variants (DMS library) of the wild-type MAGE-A3_168-176_ peptide (n = 171 variants). Heatmaps represent the proportions of CD69^high^ TnT-TCR cells after overnight co-culture, as determined by flow cytometry. Data are normalized to the response induced by wild-type peptide (boxed residues). Sequence logos show the relative activity of peptide DMS library members carrying mutations at the same peptide position. **i**, The sequences of peptides mediating 5, 10, 20, 30, 40, 50, 60, 70, 80, 90 and 100 percent activation relative to wild-type peptide were utilized to generate motifs to query the UniProtKB database. Dot plots display the number of unique human activating peptides resulting from these searches. The TnT-TCR_a3a_ cross-reactivity profile is displayed for comparison purposes. **j**, Comparison of the highest activation thresholds at which peptide DMS motifs derived from TnT-TCR_a3a_, TnT-TCR_A3-05_ and TnT-TCR_A3-10_ return off-target peptides: titin, MAGE-A6, CD166, MRCKA isoform 2 or glutamyl aminopeptidase (AMPE). The lowest activation threshold at which MAGE-A3 peptide becomes the only predicted activating antigen is also displayed. Data are displayed as mean ± SD; * *P* < 0.05, ** *P* < 0.01, *** *P* < 0.001, **** *P* < 0.0001, ns = not significant.

### Engineered TCRs expressed in primary human T cells display high target reactivity with minimal cross-reactivity

As a final validation step, we profiled the activities of selected engineered TnT-TCR_A3_ variants in primary human CD8+ T cells. To achieve this, we adapted a CRISPR-Cas9-based method for dual knockout of endogenous TCRα and TCRβ chains and simultaneous reconstitution with transgenic TCRs targeted to the TRAC locus^56,61,62^ (Fig. 5a). The lack of a complete TRAC region in the designed TCRβα HDR templates made splicing with endogenous TRAC exons a requirement for transgenic TCR surface expression. In order to confirm targeted TCR reconstitution, we first validated this approach using TCR_a3a_. Accordingly, RT-PCR using a forward TRBV5-1 (present in TCR_a3a_) primer and reverse endogenous TRAC primer yielded a 1.6 kb product only in T cells transfected with both gRNA and HDR template (Fig. 5b). Sanger sequencing of the resulting product revealed correct integration of TCR_a3a_ sequences. By contrast, a control RT-PCR amplification of TCRα chains utilizing the TRAV21 gene yielded products of expected size for test and control transfections (Fig. 5b). Flow cytometry confirmed CD3 expression and binding to MAGE-A3 peptide-MHC dextramer in T cells reconstituted with TCR_a3a_, and a greater than 95% endogenous TCR knockout efficiency (Fig. 5c).

We next generated primary human CD8+ T cells expressing wild-type TCR_A3_, TCR_a3a_, selected TCR_A3_ variants or no transgenic TCR (TCR KO). In order to verify transgenic TCR function, we first co-cultured transfected T cells with MAGE-A3-positive EJM or MAGE-A3-negative Colo205 cells, and assessed their activation by means of IFN-γ ELISpot (Figs. 5d and 5e). No significant differences in MAGE-A3-induced activation were observed in T cells transfected with wild-type TCR_A3_, TCR_A3-04_, or no TCR (Fig. 5e). Notably, while wild-type TCR_A3_ showed a minimal response to EJM cells, the complete lack of response observed in T cells transfected with TCR_A3-04_ indicated that this variant was not successfully expressed by primary T cells. In contrast, we found that T cells expressing TCR_a3a_, TCR_A3-03_, TCR_A3-05_, TCR_A3-08_ and TCR_A3-10_ were strongly and significantly activated after co-culture with EJM cells (Figs. 5d and 5e).

We next performed a series of co-culture experiments using peptide-pulsed Colo 205 cells (HLA-A*0101-positive, MAGE-A3-negative) coupled with IFN-γ ELISpot readouts. Consistent with our results using TnT-TCR cells, we observed TCR_a3a_ cross-reactivity to titin, while titin-induced responses in our engineered TCR_A3_ variants were either absent or of lower magnitude (Fig. 5f and Supplementary Fig. 18). Notably, primary T cells expressing TCR_a3a_ showed a significant titin-induced IFN-γ response relative to background, with a magnitude that was 62% of their observed MAGE-A3 response (Fig. 5f). We also observed a low but significant titin-induced IFN-γ response in primary T cells expressing TCR_A3-08_ (Fig. 5f). Importantly, we found that primary T cells expressing variants TCR_A3-03_, TCR_A3-05_ or TCR_A3-10_ displayed a significant MAGE-A3-induced IFN-γ response but undetectable activation following co-culture with titin-pulsed Colo 205 cells (Fig. 5f). Furthermore, primary T cells expressing wild-type TCR_A3_ showed a similarly negligible titin-induced response in this assay (Fig. 5f). Assessment of TCR cross-reactivity against off-target peptides previously identified for TCR_a3a_ revealed that primary T cells expressing TCR_A3-03_ or TCR_A3-08_ had significant IFN-γ responses to peptides derived from CD166, MRCKA isoform 2 and AN_HHV8 (TCR_A3-08_ only), and were thus excluded from further characterization (Supplementary Fig. 19). By contrast, primary T cells expressing TCR_A3-05_ or TCR_A3-10_ displayed high levels of specificity for the MAGE-A3 target peptide, with no other peptide inducing detectable responses.

Given the remarkable specificity of variants TCR_A3-05_ and TCR_A3-10_, we next decided to assess their cross-reactivity profiles using the TnT platform and DMS of the target MAGE-A3_168-176_ peptide (EVDPIGHLY). We assessed activation of TnT-TCR cells by flow cytometric detection of CD69 expression, as we previously validated it as a more sensitive readout for cross-reactivity detection (Fig. 4c). Similar to TnT-TCR_a3a_ peptide DMS (Fig. 4a), most mutations at positions 1, 3, 4, 5 and 9 were detrimental for TnT-TCR_A3-05_ and TnT-TCR_A3-010_ activation (Figs. 5g-h). Remarkably, we found that several peptides with mutations at positions 6 and 7, which were mostly activating in TnT-TCR_a3a_ cells, led to substantially reduced activation in TnT-TCR_A3-10_ and TnT-TCR_A3-05_ cells. Of note, we found that the presence of valine at peptide position 6, a substitution present in the titin off-target peptide (ESDPI<u>V</u>AQY), drastically reduced responses in both TnT-TCR_A3-05_ and TCR_A3-10_ (4-8% of the wild-type MAGE-A3 response), which rationalizes the lack of cross-reactivity of these engineered TCR_A3_ variants to titin (Fig. 5f). As a way of comparison, the same peptide induced a response of 91% relative to wild-type MAGE-A3 in TnT-TCR_a3a_ cells (Fig. 4a).

Querying of the UniProtKB database with motifs derived from peptide DMS data of TnT-TCR_A3-05_ and TnT-TCR_A3-10_ revealed substantial reductions in the number of predicted off-targets relative to TnT-TCR_a3a_, both in terms of human and non-human sequences (Fig. 5i and Supplementary Fig. 20). Most notably, the MAGE-A3 target peptide became the only returned human hit for TCR_A3-05_ and TCR_A3-10_ at much lower activation thresholds (26% and 44%, respectively) than TCR_a3a_ (89%), further highlighting their reduced tolerance to peptide substitutions. In a similar manner, additional human hits for TCR_A3-05_ and TCR_A3-10_ were returned at considerably lower activation thresholds relative to TCR_a3a_ (Fig. 5j).

## DISCUSSION

Here we describe the development of the TnT platform, an extensively CRISPR-edited human T cell line that supports the functional display, engineering and cross-reactivity profiling of TCRs. TnT cells harbor fully-defined genomic changes that facilitate the display and functional engineering of transgenic TCRs at high-throughput. As such, the TnT platform provides important advantages over previous TCR engineering methods that rely exclusively on affinity-based readouts, especially in light of the poor correlation that exists between TCR affinity and function^18,19^. Furthermore, TCR reconstitution by CRISPR-Cas9 targeting of the endogenous TCRβ locus offers several advantages over plasmid transfection^33^ or viral transduction^25,31,32,34–36^ such as homogenous and physiological expression of TCRs, and occurrence of a single integration event per cell ensured by targeting of the monoallelic Jurkat CDR3β genomic sequence. Furthermore, optimization of our CRISPR-Cas9 HDR protocol led to TCR integration efficiencies of up to 20%, which allowed for the screening of large mutagenesis libraries and selection of variants with enhanced specificity and function.

The unique features of the TnT platform allowed us to apply CDR3β DMS to comprehensively profile two tumor-reactive TCRs (TCR_A3_ and TCR_DMF4_) for their patterns of expression, antigen binding and antigen-induced signaling. Deep sequencing of original libraries and FACS selections was a crucial component of our TCR engineering pipeline (both in DMS and combinatorial selections) as it allowed us to accurately determine the enrichment of specific TCR variants across selections. Remarkably, we identified several variants that were enriched for antigen binding but not for antigen-induced signaling, and vice-versa, which emphasized the discordance between TCR binding and function. For example, two TCR_DMF4_ variants had nearly identical signaling capacity despite one of them displaying undetectable binding to the MART-1 peptide-MHC dextramer. This is in agreement with previous reports of peptide-MHC multimers failing to detect fully functional TCRs^42^. Crucially, DMS enrichment data allowed us to tailor the design of a combinatorial TCR_A3_ library in order to maximize the number of productive variants.

We utilized the TnT platform for engineering the MAGE-A3-specific TCR_A3_, a naturally occurring low-avidity TCR that has been previously engineered by phage display, which resulted in affinity-enhanced variants with cross-reactivity to titin^14,30,63^. To this end, we designed a DMS-based combinatorial library of CDR3β TCR_A3_ and used the TnT platform to perform selections based on binding and signaling in response to both MAGE-A3 and titin antigens. Throughout the selection process, we observed the emergence of TCR_A3_ variants with enhanced binding and signaling in response to MAGE-A3. Interestingly, we found that a considerable proportion of library members also displayed binding and signaling in response to titin, highlighting the high propensity of TCR_A3_ mutants to develop titin cross-reactivity. By using deep sequencing enrichment data, we could identify TCR_A3_ variants with increased binding and function in response to MAGE-A3 that neither bound or responded to titin. Surprisingly, we found that some variants (e.g., TCR_A3-27_ and TCR_A3-28_) displayed robust signaling in response to titin, despite showing low or undetectable binding to titin-MHC dextramer. This finding further emphasizes the importance of functional screening of engineered TCRs, as screening approaches based on binding alone may fail to identify cross-reactive TCRs^16,40^. Remarkably, TCR_A3-05_, which showed the highest specificity to MAGE-A3 throughout our experiments in both TnT cells and primary T cells, was the variant with the highest score for predicted MAGE-A3 specificity based on deep sequencing enrichment data (Supplementary Table 3). Taken together, these findings highlight the power of TnT functional selections coupled with deep sequencing for engineering highly specific TCRs.

The TnT platform allows us to both engineer TCRs and assess their cross-reactivity potential without the need of reformatting TCRs for cellular display^31,48,49^ or soluble expression^39,40^. By performing DMS of the target MAGE-A3_168-176_ peptide, we predicted and validated known and potentially novel off-targets for TCR_a3a_. Despite its four amino acid difference to MAGE-A3, the titin ESDPIVAQY peptide was identified at the second highest activation threshold. Interestingly, we also identified a peptide derived from MRCKA isoform 2 (UniProtKB, Q5VT25-2) as a potential TCR_a3a_ off-target, and confirmed its ability to activate TnT-TCR_a3a_ cells *in vitro*. As MRCKA is highly expressed in the heart^64^, it is possible that recognition of this peptide may have also contributed to the cardiac toxicity elicited by TCR_a3a_^14^. We observed that a number of our engineered TCR_A3_ variants were activated by predicted TCR_a3a_ off-targets. As these TnT-TCR_A3_ variants showed weaker binding to MAGE-A3 peptide-MHC than TnT-TCR_a3a_ (TCR_a3a_ *K_D_* ~ 2.3 – 6.6 μM)^23,30^, they constitute examples of cross-reactivity emergence in engineered TCRs within the physiological affinity range (1 – 100 μM)^19^.

Assessment of primary T cell activation by means of IFN-γ ELISpot assays showed a higher sensitivity than CD69 expression in TnT-TCR cells following co-culture, which may reflect higher levels of co-receptors in primary CD8+ T cells relative to TnT cells. Despite this difference, we were able to identify variants that displayed remarkable specificity for MAGE-A3, namely TCR_A3-05_ and TCR_A3-10_, in both TnT-TCR and primary T cell assays. Functional cross-reactivity profiling of these variants using the TnT platform revealed a large reduction in the number of predicted off-targets compared to TCR_a3a_. While reduced cross-reactivity was particularly evident for titin, it was also observed for peptides that were not negatively selected for (e.g., MAGE-A6, CD166 and MRCKA isoform 2), thus highlighting TCR_A3-05_ and TCR_A3-10_ as intrinsically specific variants. This is consistent with the recent identification of naturally-occurring TCRs displaying a wide range of specificity levels for the same target^16^. Thus, the enhanced function and specificity of TCR_A3-05_ and TCR_A3-10_ make them promising candidates for use in TCR gene therapies targeting MAGE-A3-positive tumors in HLA-A*0101-positive patients. Thus, the ability of the TnT platform to support both TCR engineering and the accurate prediction of cross-reactivity on the basis of function makes it a promising technology for the development of therapeutic TCRs with improved efficacy and safety profiles.

## METHODS

### Cell lines and cell culture

The Jurkat leukemia E6-1 T cell line was obtained from the American Type Culture Collection (ATCC) (#TIB152); the T2 hybrid cell line (#ACC598) and the EJM multiple myeloma cell line (#ACC560) were obtained from the German Collection of Cell Culture and microorganisms (DSMZ); and the Colo 205 colon adenocarcinoma cell line (#87061208) was obtained from the European Collection of Authenticated Cell Cultures (ECACC). Jurkat cells, engineered TnT cells and Colo 205 cells were cultured in ATCC-modified RPMI-1640 (Thermo Fisher, #A1049101), T2 cells were cultured in RPMI-1640 (Thermo Fisher, #11875093), and EJM cells were cultured in IMDM (Thermo Fisher, #12440053). All media were supplemented with 10% FBS, 50 U ml^−1^ penicillin and 50 μg ml^−1^ streptomycin. Detachment of EJM and Colo 205 adherent cell lines for passaging was performed using the TrypLE reagent (Thermo Fisher, #12605010). All cell lines were cultured at 37 °C, 5% CO_2_ in a humidified atmosphere.

### Polymerase chain reaction (PCR)

PCRs for cloning, generation of HDR templates, genotyping of mammalian cells, generation of TCR libraries and generation of amplicons for deep sequencing were performed using the KAPA HiFi PCR kit with GC buffer (Roche Diagnostics, #07958846001) and custom designed primers (Supplementary Table 6). Annealing temperatures (x) were optimized for each reaction by gradient PCR and cycling conditions were as follows: 95°C for 3 min; 35 cycles of 98°C for 20 s, x°C for 15 s, 72°C for 30 s per kb; final extension 72°C for 1 min per kb. PCRs for genotyping of bacterial colonies after transformation were performed using the KAPA2G Fast ReadyMix kit (Sigma Aldrich, #KK5102) with custom designed primers and the following cycling conditions: 95°C for 3 min; 35 cycles of 95°C for 15 s, 60°C for 15 s, 72°C for 15 s per kb; final extension 72°C for 1 min per kb.

### Cloning and generation of HDR templates

DNA for gene-encoding regions and homology regions were generated by gene synthesis (Twist Bioscience) or PCR and introduced into desired plasmid backbones via restriction cloning (Supplementary File 1). The following plasmids were used as backbones: pX458 (Addgene, #48138), AAVS1_Puro_Tet3G_3xFLAG_Twin_Strep (Addgene, #92099), pGL4.30 (Promega, #E8481) and pTwist Amp High Copy (Twist Bioscience). Targeted knock-in of Cas9/GFP into the CCR5 locus was performed utilizing circular plasmid DNA as the HDR template. HDR templates for all other targeted knock-in experiments were provided as linear double-stranded DNA (dsDNA) generated by PCR. Prior to transfection, PCR products were column-purified using the QIAquick PCR Purification Kit (Qiagen, #28106). For targeted TCR reconstitution of TnT cells, homology arms flanking the recombined Jurkat TCRβ VDJ locus were designed and cloned in pTwist (Twist Bioscience), resulting in pJurTCRB. TCRαβ cassettes encoding transgenic TCRs were generated by gene synthesis (Twist Bioscience) and cloned into pJurTCRB using naturally-occurring XbaI and BsaI restriction sites present within the homology arms. Next, HDR templates were generated by PCR using primer pair RVL-127/128 and PCR products purified prior to transfection. For targeted TCR reconstitution of primary human CD8+ T cells, TCRβα cassettes lacking TRAC exons 2-3 and flanked by homology arms mapping to TRAC exon 1^56^ were designed and cloned in pTwist (Twist Bioscience). HDR templates were generated by PCR using primer pair RVL-164/165 and PCR products purified prior to transfection.

### CRISPR-Cas9 genome editing

Transfection of TnT cells and Jurkat-derived cell lines was performed by electroporation using the 4D-Nucleofector device (Lonza) and the SE cell line kit (Lonza, #V4XC-1024). The day before transfection, cells were seeded at 2.5×10^5^ cells/mL and cultured for 24 h. Prior to transfection, gRNA molecules (Supplementary Table 7) were assembled by mixing 4 μl of custom Alt-R crRNA (200 μM, IDT) with 4 μL of Alt-R tracrRNA (200 μM, IDT, #1072534), incubating the mix at 95°C for 5 min and cooling it to room temperature. For transfection of Cas9-negative cell lines, 2 μL of assembled gRNA molecules were mixed with 2 μL of recombinant SpCas9 (61 μM, IDT, #1081059) and incubated for > 10 min at room temperature to generate Cas9 RNP complexes. Immediately prior to transfection, cells were washed twice in PBS and 1×10^6^ cells were re-suspended in 100 μL of SE buffer. 1.5 μg of HDR template and 7 μL of assembled gRNA (or 4 μL of Cas9 RNP complexes) were added to the cell suspension, mixed and transferred into a 1 mL electroporation cuvette. Cells were electroporated using program CK116, topped-up with 1 mL of complete media and rested for 10 min prior to transfer into a 12-well plate. Alt-R HDR enhancer (IDT, #1081073) was added at a 30 μM final concentration and removed after 16 h of culture by centrifugation. HDR efficiency was assessed by flow cytometry on day 5 post-transfection. For transfections at the 20 μL scale (Lonza, #V4XC-1032), cell numbers and reagent volumes were reduced 5-fold.

### Flow cytometry and fluorescence-activated cell sorting (FACS)

Flow cytometric analysis of cell lines and primary T cells was performed according to standard protocols. The following antibodies were purchased from BioLegend and used at 1 μg ml^−1^ in flow cytometry buffer (PBS, 2% FBS, 2 mM EDTA): PE-Cy7-conjugated or APC-conjugated anti-human CD3e (clone UCHT1, #300420 or #300458), APC-conjugated anti-human CD4 (clone RPA-T4, #300552), PE-conjugated anti-human CD8a (clone HIT8a, #300908), PE-Cy7-conjugated anti-human CD19 (clone HIB19, #302216), APC-conjugated anti-human CD69 (clone FN50, #310910), APC-conjugated anti-human Fas (clone DX2, #305611) and PE-conjugated anti-human TCR α/β (clone IP26, #306707). DAPI viability dye (Thermo Fisher, #62248) was added to antibody cocktails at a final concentration of 1 μg ml^−1^. Cells were washed once in flow cytometry buffer prior to staining, stained for 20 min on ice and washed twice in flow cytometry buffer before analysis using BD LSRFortessa or Beckman-Coulter CytoFLEX flow cytometers. Blocking of Fc receptors in T2 cells was performed prior to staining using the TruStain FcX reagent (BioLegend, #422301). Staining with peptide-MHC dextramers was performed for 10 min at room temperature (RT), followed by addition of 2X antibody cocktails (2 ug ml^−1^ antibodies, 2 ug ml^−1^ DAPI) and incubation for 20 min on ice. The following peptide-MHC dextramers were commercially obtained from Immudex: NY-ESO-1_157-165_ (SLLMWITQC, HLA-A*0201, #WB2696-APC); MART-1_26-35(27L)_ (ELAGIGILTV, HLA-A*0201, #WB2162-APC); MAGE-A3_168-176_ (EVDPIGHLY, HLA-A*0101, #WA3249-PE) and titin_24,337-24,345_ (ESDPIVAQY, HLA-A*0101, custom-made, APC-conjugated). Peptide-MHC dextramers were used at a 3.2 nM final concentration (i.e., 1:10 dilution) for staining, unless indicated otherwise in figure legends. FACS was performed using BD FACSAria III or BD FACSAria Fusion instruments. Single-cell sorts were collected in 96-well flat-bottom plates containing conditioned media and clones were cultured for 2-3 weeks prior to characterization.

### Genotyping of cell lines and transfectants

Genomic DNA was extracted from 2×10^5^ cells by resuspension in 100 μL of QuickExtract solution (Lucigen, #0905T), incubation at 65°C for 6 min, vortexing for 15 s and incubation at 98°C for 2 min. 5 μL of genomic DNA extract were then used as templates for 25 μL PCR reactions. For genotyping by two-step reverse transcription PCR (RT-PCR), RNA from 1×10^5^ cells was extracted using the TRIZol reagent (Invitrogen, # 15596018) and column-purified using the PureLink RNA Mini kit (Invitrogen, #12183025). For reverse transcription, 100 pmol of oligo dT, 10 nmol of each dNTP, 5 μL RNA and sufficient nuclease-free water for a final 14 μl volume were mixed, incubated at 65°C for 5 min and chilled on ice for 5 min. This was followed by addition of 4 μL of 5X RT buffer, 40 units of RiboLock RNAse inhibitor (Thermo Fisher, #EO0381) and 200 units of Maxima H-minus reverse transcriptase (Thermo Fisher, #EP0751) and mixing. In some experiments, 40 pmol of template-switching oligonucleotide (TSO, Supplementary Table 6) was added for labelling of first-strand cDNA 3’ ends^65^. Reverse transcription was performed at 50°C for 30 min, followed by inactivation at 85°C for 5 min. 5 μl of the resulting cDNA-containing reverse transcription mixes were then used as templates for 25 μL PCR reactions.

### Peptides and peptide pulse

Peptides and peptide libraries were generated by custom peptide synthesis (Genscript), re-suspended at 10 mg ml^−1^ in DMSO and placed at −80°C for prolonged storage. For peptide pulsing, T2 cells or Colo 205 cells were harvested and washed twice in serum-free RPMI 1640 (SF-RPMI). Peptides were diluted to 10 μg ml^−1^ in SF-RPMI (or to concentrations indicated in figure legends) and the solution was used to re-suspend cells at 1×10^6^ cells ml^−1^. Cells were incubated for 90 min at 37°C, 5% CO2, washed once with SF-RPMI, re-suspended in complete media and added to co-culture wells (see section below).

### TnT stimulation and co-culture assays

For clone screening and assessment of AICD, TnT cells and Jurkat-derived cell lines were stimulated overnight with either 10 μg ml^−1^ plate-bound anti-human CD3e antibody (clone OKT3, BioLegend, #317326) or 1X eBioscience Cell Stimulation Cocktail (81 nM PMA, 1.34 μM ionomycin; Thermo Fisher, #00497093). For co-culture experiments, TnT-TCR cells at ~ 1×10^6^ cells ml^−1^ density were harvested, pelleted by centrifugation and re-suspended in fresh complete media at 1×10^6^ cells ml^−1^. 1×10^5^ TnT-TCR cells (100 μL) were seeded in wells of a V-bottom 96-well plate. Antigen-expressing cells (EJM) or peptide-pulsed cells (T2, Colo 205) were adjusted to 1×10^6^ cells ml^−1^ in complete media and 5×10^4^ cells (50 μL) added to each well. Anti-human CD28 antibody (clone CD28.2, BioLegend, #302933) was added at a final concentration of 1 μg ml^−1^ for co-stimulation of all samples (including negative controls) and plates were incubated overnight at 37°C, 5% CO_2_. The next day, expression of NFAT-GFP and CD69 in TnT-TCR cells was assessed by flow cytometry. Flow cytometric discrimination between TnT-TCR cells and Colo 205 cells (or EJM cells) was based on side scatter area (SSC-A) and mRuby expression, while discrimination between TnT-TCR cells (CD19-negative) and T2 cells (CD19-positive) was based on CD19 expression.

### Generation of deep mutational scanning (DMS) libraries

DMS libraries of the CDR3β regions of TCR_A3_ and TCR_DMF4_ were generated by plasmid nicking mutagenesis as described in Wrenbeck et al. 2016^59^. The protocol relies on the presence of a single BbvCI restriction site for sequential targeting with Nt.BbvCI and Nb.BbvCI nickases, digestion of wild-type plasmid and plasmid re-synthesis using mutagenic oligonucleotides. A plus-strand BbvCI restriction site was introduced into the pJurTCRB-TCR_A3_ plasmid by means of PCR and blunt-end ligation, while the endogenous minus-strand BbvCI site present in the TRBV10-3 gene of pJutTCRB-TCR_DMF4_ was targeted. The order of BbvCI nickase digestion was adjusted for each plasmid so that the plus DNA strand was digested first. Mutagenic oligonucleotides were designed using the QuikChange Primer Design online tool (Agilent) and assessed for the presence of secondary structures using the Oligo Evaluator online tool (Sigma-Aldrich) (Supplementary Table 6). Oligonucleotides showing strong potential for forming secondary structures were manually modified to reduce this propensity. After nicking mutagenesis, mutated plasmids were transformed into 100 μL of chemically-competent *E. coli* DH5α cells (NEB, #C2987H) and plated on ampicillin (100 μg ml^−1^) LB agar in Nunc BioAssay dishes (Sigma-Aldrich, #D4803). Serial dilutions of transformed cells were plated separately to quantify bacterial transformants. Plasmid libraries were purified from bacterial transformants using the QIAprep Spin Miniprep kit (Qiagen, #27106). HDR templates were generated from plasmid libraries by PCR using primer pair RVL-127/128 and column-purified prior to transfection.

### DMS library screening and selections

DMS library HDR templates and CDR3B gRNA were used to transfect 1×10^6^ TnT cells. In TCR_A3_ DMS selections, cells with restored CD3 surface expression and no binding to control titin peptide-MHC dextramer were isolated by FACS on day 8 post-transfection (SEL 1). Sorted cells were expanded for 13 days and either stained with MAGE-A3 peptide-MHC dextramer or co-cultured overnight with MAGE-A3-positive EJM cells. Dextramer-positive cells (SEL 2A) and activated CD69^high^ cells (SEL 2B) were then isolated by FACS. In TCR_DMF4_ DMS selections, cells with restored CD3 surface expression and no binding to control NY-ESO-1 peptide-MHC dextramer were isolated by FACS on day 8 post-transfection (SEL 1). Sorted cells were expanded for 13 days and either stained with MART-1 peptide-MHC dextramer or co-cultured overnight with MART-1 peptide--pulsed T2 cells. Dextramer-positive cells (SEL 2A) and activated NFAT-GFP-positive cells (SEL 2B) were then isolated by FACS.

### Generation of combinatorial TCR_A3_ libraries

Degenerate codons reflecting the combined CDR3β amino acid frequencies observed in TCR_A3_ DMS binding and signaling selections (SEL2A+2B) were determined as previously described^51^. The library resulting from two iterations of our algorithm was modified to include VNB codons at CDR3β positions 4 and 6. For library construction, ssDNA oligonucleotides containing a 28 nt complementary overlap were designed and purchased as custom ultramers (IDT, Supplementary Table 6). The forward ultramer encoded exclusively wild-type TCR_A3_ codons, while the reverse ultramer contained the reverse complement of both wild-type and library degenerate codons. 200 pmol of each ultramer were mixed and subjected to single-cycle PCR using the following conditions: 95°C for 3 min, 98°C for 20 s, 70°C for 15 s, 72°C for 10 min. The resulting 270 bp dsDNA product was gel-purified (Zymogen, #D4002) and 8 ng were utilized as template for a 200 μL PCR reaction using external primers with the following cycling conditions: 95°C for 3 min; 25 cycles of 98°C for 20 s, 62°C for 15 s, 72°C for 15 s; final extension 72°C for 30 s. The PCR product was column-purified, digested with KpnI and BsaI restriction enzymes, and re-purified. In parallel, the pJurTCRB-TCR_A3_ plasmid was digested with KpnI and BsaI, de-phoshorylated (CIP, NEB, #M0290) and gel-purified. Digested PCR product (112.5 ng) and plasmid (750 ng) were ligated in a 75 μl reaction containing 1X T4 PNK buffer, 1 mM ATP and 3 units of T4 DNA ligase for 2 h at RT (all from NEB). Next, the ligation mix was transformed into 750 μL of chemically-competent *E. coli* DH5α cells (NEB, #C2987I) and plated on ampicillin LB agar in Nunc BioAssay dishes. Quantification of bacterial transformants, purification of plasmid library and generation of HDR templates was performed as described for DMS libraries.

### Combinatorial TCR_A3_ library screening and selections

Combinatorial library HDR templates (20 μg) and CDR3B gRNA (10 nmol) were used to transfect 1×10^8^ TnT cells using the 4D-Nucleofector LV unit (Lonza, #AAF-1002L). TnT cells with restored CD3 surface expression were bulk-sorted (SEL 1) on day 6 post transfection. SEL 1 cells were expanded for 6 days prior to overnight co-culture with MAGE-A3-positive EJM cells followed by co-staining with MAGE-A3 and titin peptide-MHC dextramers. After co-culture, NFAT-GFP-positive cells displaying positive MAGE-A3 peptide-MHC binding and negative titin peptide-MHC binding were bulk-sorted (SEL 2) and expanded in culture for 12 days. SEL 2 cells were co-cultured overnight with either peptide MAGE-A3-pulsed (MAGE-A3) or titin-pulsed Colo 205 cells. Activated NFAT-GFP-positive cells from MAGE-A3 (SEL 3A) and titin (SEL 3B) co-cultures were bulk-sorted for RNA extraction, RT-PCR and deep sequencing.

### Deep sequencing and analysis of TCR libraries

TCR amplicons for deep sequencing of plasmid libraries were generated by PCR using primer pair RVL-144/154, while TCR amplicons for deep sequencing of TnT-TCR selections were generated by two-step RT-PCR using primer pair RVL-144/145. In both cases, PCR was limited to 25 cycles. TCR amplicons were column-purified and deep-sequenced using the Amplicon-EZ service (Genewiz), which includes adaptor/index ligation and paired-end Illumina sequencing (250 cycles) followed by delivery of 50,000 assembled reads per sample with unique sequence identification and abundance analysis. For DMS plasmid libraries and selections, unique sequences with less than ten sequencing reads were excluded from enrichment analysis, as every library member had sequencing reads above this threshold. Sequence enrichment of unique DMS variants was determined by dividing their observed frequencies in SEL 1 (TCR-CD3 expression), SEL 2A (binding) and SEL 2B (signaling) over their plasmid DMS library frequencies, and heatmaps were generated using the GraphPad Prism software. For the TCR_A3_ combinatorial plasmid library and selections, unique clone frequency data was filtered to remove clones containing insertions, deletions or mutations outside CDR3β. Filtered data was used to generate sequence logos weighted on amino acid frequencies at specific CDR3β positions using R packages ggseqlogo^66^ and ggplot2. The frequencies of specific TCR_A3_ variants across selections were identified by merging unique clone datasets using a custom Python script. Sequence enrichment of unique TCR_A3_ combinatorial variants was determined by dividing their observed frequencies in SEL 2 (MAGE-A3-induced activation and binding), SEL 3A (MAGE-A3-induced activation) and SEL 3B (titin-induced activation) over their SEL 1 (TCR-CD3 expression) frequencies.

### Peptide DMS and assessment of TCR cross-reactivity

A DMS library of the target MAGE-A3_168-176_ EVDPIGHLY peptide was designed and generated by custom peptide synthesis (Genscript). Each library member (n = 171) was individually pulsed at a 50 ug ml^−1^ concentration on Colo 205 cells for co-culture with TnT-TCR cells (n = 171 co-cultures). Co-cultures with MAGE-A3-pulsed (n = 3), titin-pulsed (n = 3), CMV-pulsed (n = 6) peptides and unpulsed (n = 6) Colo 205 cells were included as controls. After overnight co-culture, TnT-TCR activation was assessed by NFAT-GFP and CD69 expression by means of flow cytometry. The mean background activation observed in CMV peptide controls (i.e., VTEHDTLLY peptide pulse) was subtracted from all samples and their responses normalized to the mean MAGE-A3 response level. Normalized data was used to generate heatmaps (GraphPad Prism), weighted sequence logos (ggseqplot, ggplot2 in R) and peptide sequence motifs of allowed substitutions at discrete activation thresholds (Bioconductor package Biostrings in R). Peptide sequence motifs (Supplementary Table 4) were then used to query the UniProtKB database (including splice variants) with the ScanProsite online tool. The output of these searches was processed using the Biostrings package in order to compute the number of unique peptide hits.

### Primary T cell culture and genome editing

Human peripheral blood mononuclear cells were purchased from Stemcell Technologies (#70025) and CD8+ T cells isolated using the EasySep Human CD8+ T Cell Isolation kit (Stemcell Technologies, #17953). Primary human CD8+ T cells were cultured for up to 24 days in ATCC-modified RPMI (Thermo Fisher, #A1049101) supplemented with 10% FBS, 10 mM non-essential amino acids, 50 μM 2-mercaptoethanol, 50 U ml^−1^ penicillin, 50 μg ml^−1^ streptomycin and freshly added 20 ng ml^−1^ recombinant human IL-2, (Peprotech, #200-02). T cells were activated with anti-CD3/anti-CD28 tetrameric antibody complexes (Stemcell Technologies, #10971) on days 1 and 13 of culture and expanded every 3-4 days. Transfection of primary T cells with Cas9 RNP complexes and TCRβα HDR templates was performed 3-4 days following activation using the 4D-Nucleofector and a 20 uL format P3 Primary Cell kit (Lonza, V4XP-3032). Briefly, 1×10^6^ primary CD8+ T cells were transfected with 1 μg of HDR template, 1 μl of TRAC Cas9 RNP complex and 1 μl of TRBC1/2 Cas9 RNP complex using the EO115 electroporation program (Cas9 RNP complexes = 50 μM gRNA, 30.5 μM recombinant SpCas9). For RT-PCR validation of TCR reconstitution, RNA was extracted from 1×10^6^ T cells, quantified using a Nanodrop instrument, and 40 ng RNA used as input for reverse transcription. 2 μL of reverse transcription mixes were then utilized as templates for 25 μL PCR reactions.

### Co-culture of primary T cells and IFN-γ ELISpot

IFN-γ ELISpot assays were performed using the Human IFN-γ ELISpot Pair (BD, #551873), 96-well ELISpot plates (Millipore, #MSIPS4W10), Avidin-HRP (Biolegend, #405103) and precipitating TMB substrate (Mabtech, #3651-10). Wells were activated with 15%(v/v) ethanol for 30 s, washed twice with PBS and coated with 5 μg ml^−1^ capture antibody (in PBS) at 4°C overnight (or up to a week). On the day of co-culture (i.e, day 5 post-transfection), wells were washed twice with PBS and blocked with primary T cell media lacking IL-2 (RP10-TC) for > 2 h at 37°C. In parallel, TCR-reconstituted primary CD8+ T cells were rested in the absence IL-2 for 6 h. After resting, T cells were washed and re-suspended in fresh RP10-TC media. A 100 μL volume of cell suspensions containing 5×10^4^ to 4×10^5^ T cells was then transferred into blocked ELISpot wells, as specified in figure legends. Next, 1.5×10^4^ antigen-expressing (EJM) or peptide-pulsed (Colo 205) cells were added into wells in a 50 μL volume of RP10-TC media. Anti-CD28 monoclonal antibody was added into every well at a 1 μg ml^−1^ final concentration and plates were incubated for 20 h at 37°C, 5% CO_2_. Following co-culture, cells were removed and wells washed three times with wash buffer (0.01%(v/v) Tween 20 in PBS). Detection antibody was then added at 2 μg ml^−1^ in dilution buffer (0.5%(v/v) BSA in PBS) followed by 2 h incubation at RT. After incubation, wells were washed three times with wash buffer and 1:2000 avidin-HRP (in dilution buffer) added for 45 min at RT. Wells were washed three times with wash buffer and once in PBS, followed by development with precipitating TMB substrate for 3-10 min at RT. Development was stopped by washing with deionized water and plates were dried for > 24 h in the dark prior to analysis using an AID ELR08 ELISpot reader (Autoimmun Diagnostika).

## AUTHOR CONTRIBUTIONS

R.V.-L. and S.T.R. designed the study; R.V.-L., J.J. and S.T.R. contributed to experimental design; R.V.- L., J.J., F.B., E.A. and E.K. performed experiments; R.V.-L., J.J., F.B., and C.R.W. analyzed data; R.V.-L. and S.T.R. wrote the manuscript with input from all authors.

## ACKNOWLEDGMENTS

We thank Dr. Joseph Taft for his advice on the plasmid nicking mutagenesis method, and Darya Palianina for her assistance with ELISpot assays. We acknowledge the ETH Zurich D-BSSE Single Cell Unit for their assistance with FACS. This study is supported by funding from the Personalized Health and Related Technologies Postdoctoral Fellowship (to R.V.-L), the NCCR Molecular Systems Engineering (to S.T.R.) and Helmut Horten Stiftung (to S.T.R.).

## DATA AVAILABILITY

The raw FASTQ files from deep sequencing that support the findings of this study will be deposited in the Sequence Read Archive (SRA) with the primary accession code(s) <code(s) (https://www.ncbi.nlm.nih.gov/sra)>. Additional data that support the findings of this study are available from the corresponding author upon reasonable request.

