## Supplementary Material for "CRISPR-targeted display of functional T cell receptors enables engineering of enhanced specificity and prediction of cross-reactivity"

[1]To whom correspondence should be addressed

Supplementary Figures 1-20

Supplementary Tables 1-7

### Supplementary Figures 1-20

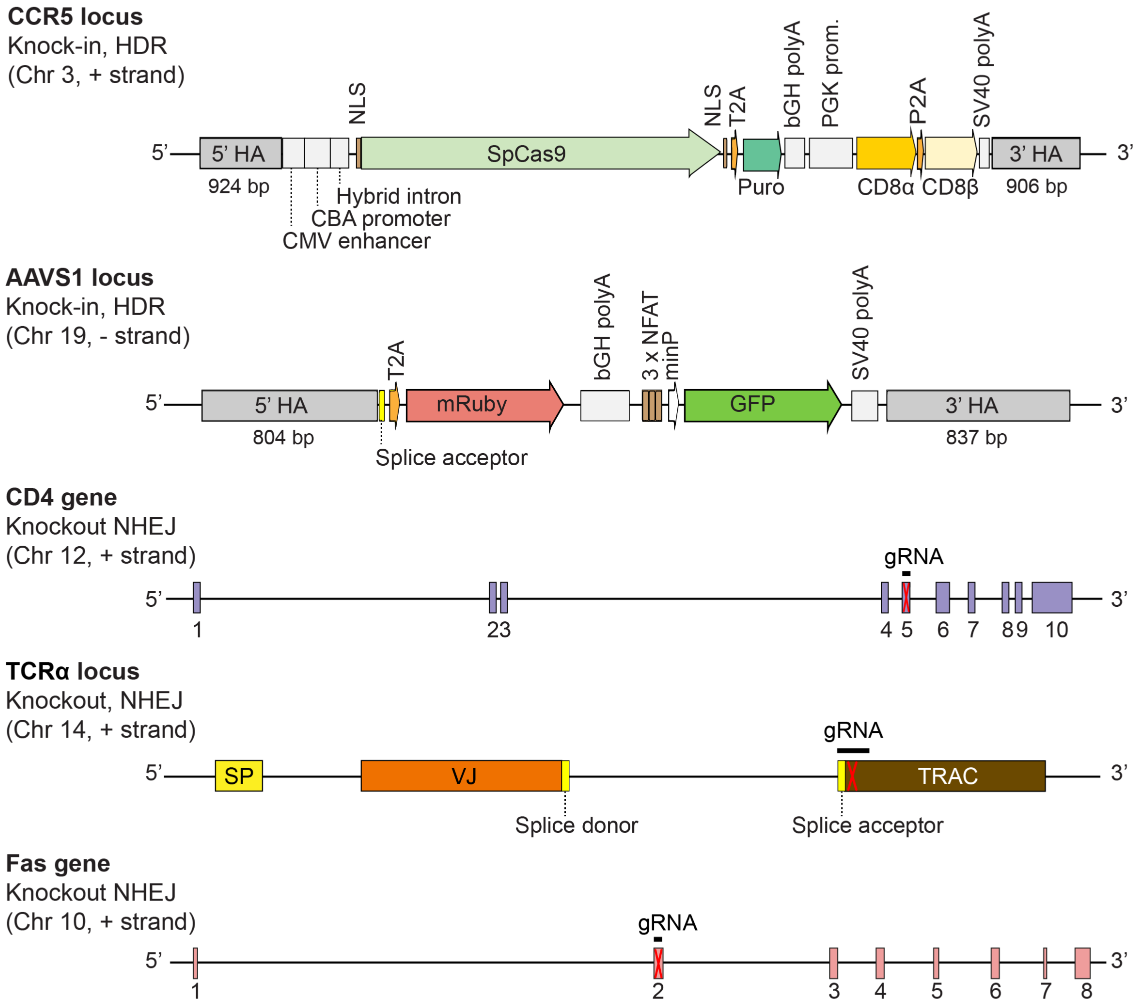


**Supplementary Figure 1. Development of the TCR-accepting T cell (TnT) platform through multistep CRISPR-Cas9 genome editing.** Schematic representation of the location and changes of edited genomic loci in the TnT platform. AAVS1: adeno-associated virus integration site 1, bGH polyA: bovine growth hormone polyadenylation signal, GFP: green fluorescent protein, gRNA: guide RNA, HA: homology arm, HDR: homology-directed repair, minP: minimal promoter, NHEJ: non-homologous end joining, NLS: nuclear localization signal, P2A: 2A peptide from porcine teschovirus-1 polyprotein, PGK: phosphoglycerate kinase. Puro: puromycin, SP: signal peptide, SpCas9: *Streptococcus pyogenes* Cas9, T2A: 2A peptide from *Thosea asigna* virus capsid protein, TRAC: T cell receptor alpha constant region, SV40 polyA: simian virus 40 polyadenylation signal.

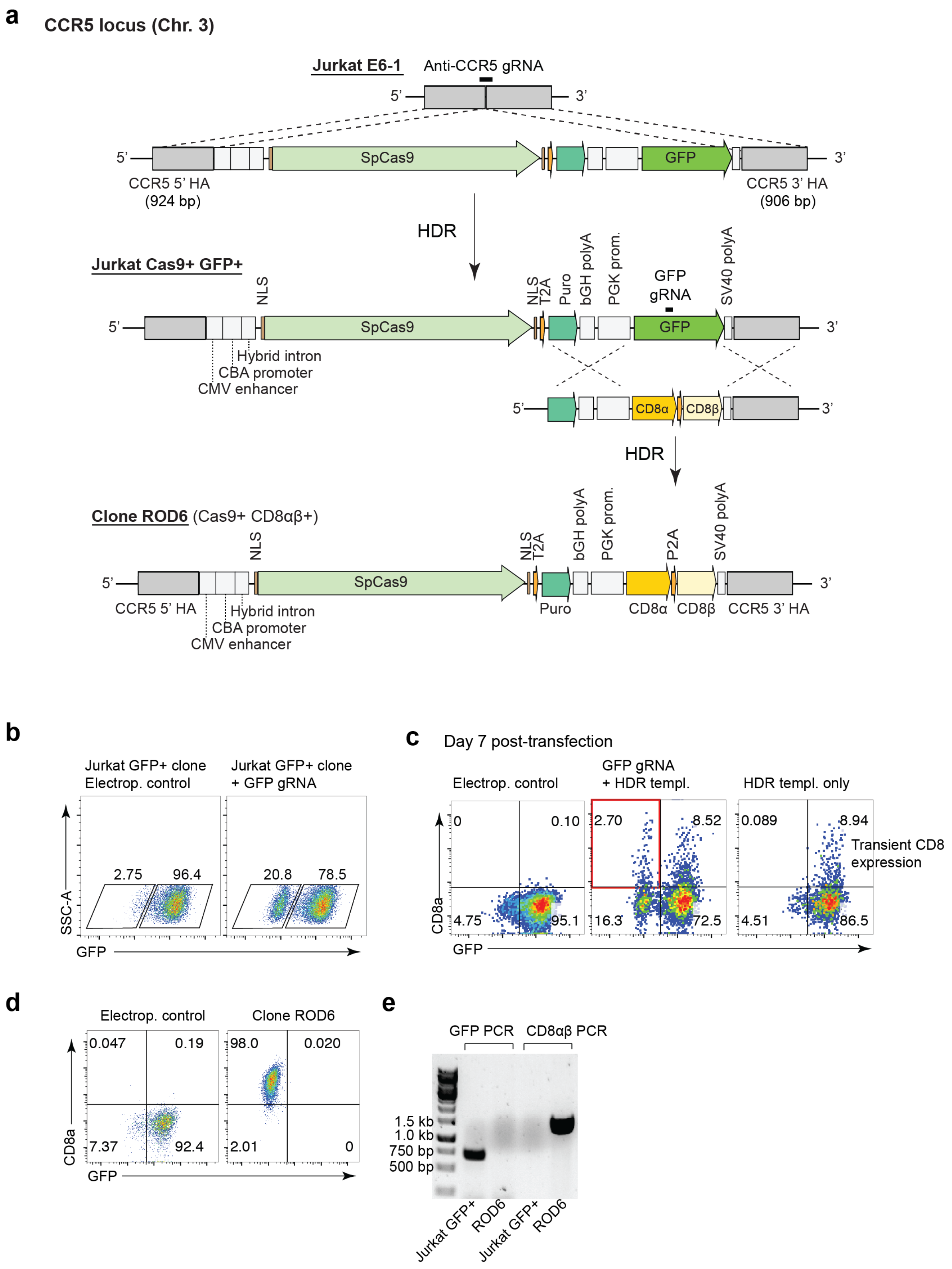
a
CCR5 locus (Chr. 3)
Jurkat E6-1
Anti-CCR5 gRNA
SpCas9
GFP
CCR5 5' HA (924 bp)
CCR5 3' HA (906 bp)
HDR
Jurkat Cas9+ GFP+
NLS
T2A
Puro
bGH polyA
PGK prom.
GFP gRNA
SV40 polyA
Hybrid intron
CBA promoter
CMV enhancer
CD8α
CD8β
Clone ROD6 (Cas9+ CD8αβ+)
P2A
CCR5 5' HA
CCR5 3' HA
b
Jurkat GFP+ clone Electrop. control
Jurkat GFP+ clone + GFP gRNA
2.75
96.4
20.8
78.5
SSC-A
GFP
c
Day 7 post-transfection
Electrop. control
GFP gRNA + HDR templ.
HDR templ. only
0
0.10
4.75
95.1
2.70
8.52
16.3
72.5
0.089
8.94
4.51
86.5
Transient CD8 expression
CD8a
d
Electrop. control
Clone ROD6
0.047
0.19
7.37
92.4
98.0
0.020
2.01
0
e
GFP PCR
CD8αβ PCR
1.5 kb
1.0 kb
750 bp
500 bp
Jurkat GFP+
ROD6
Jurkat GFP+
ROD6

**Supplementary Figure 2. CRISPR-Cas9 integration of Cas9 and human CD8 into the CCR5 safe harbour-locus of Jurkat T cells. a,** Schematic representation of the two-step approach utilized for the introduction of Cas9 and CD8αβ genes into the CCR5 locus. The CCR5 genomic loci in the parental Jurkat E6-1 cell line, intermediate Jurkat Cas9+ GFP+ cell line and final Jurkat cell line (ROD6 clone) are displayed. **b,** A Jurkat GFP+ clone was initially derived by transfecting Jurkat E6-1 cells with an CCR5 gRNA complexed with recombinant Cas9 protein (IDT) and a plasmid HDR template consisting of CCR5 homology arms and genes encoding *S. pyogenes* Cas9, puromycin N-acetyl transferase and green fluorescent protein (GFP). Transfection of the selected Jurkat GFP+ clone with GFP gRNA alone (i.e., no external source of Cas9) led to GFP knockout, thus demonstrating activity of the endogenously expressed Cas9 in this cell line. **c,** The introduced GFP gene was targeted for replacement with human CD8 by means of CRISPR-Cas9 genome editing. An HDR template with homology arms mapping to the transgenic puromycin resistance gene (5') and the CCR5 genomic locus (3'), and flanking a construct encoding hCD8α-P2A-hCD8β was designed. Flow cytometry shows results of transfection of Jurkat Cas9+ GFP+ cells with the HDR template above (PCR product) and a GFP gRNA, yielding the desired GFP- CD8α+ cells (red box). **d,** GFP- CD8α+ cells in (c) were subjected to single-cell FACS for the isolation of clone ROD6. **e,** PCR amplification of GFP and CD8α-P2A-CD8β genes from genomic DNA of clone ROD6. Subsequent Sanger sequencing verified the successful replacement of GFP with human CD8.

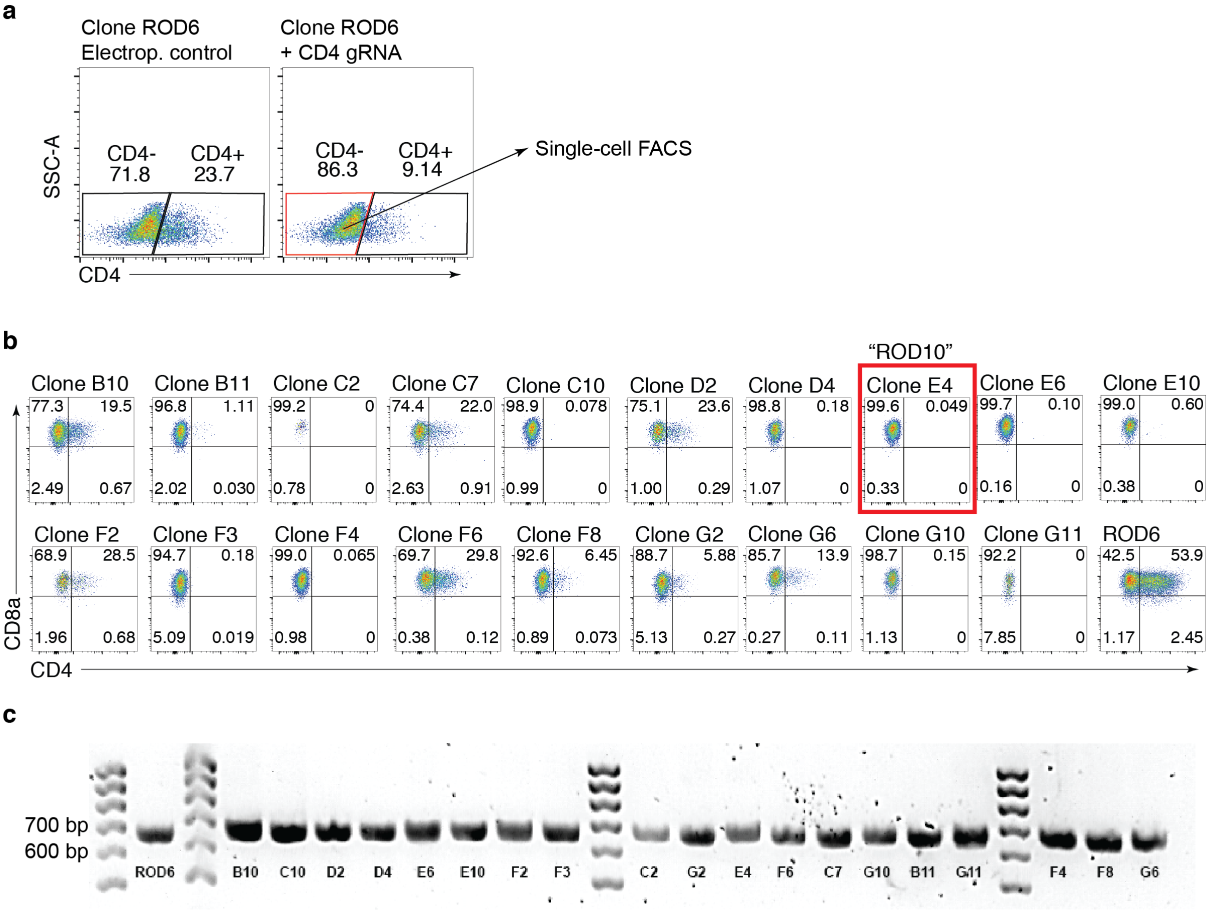


**Supplementary Figure 3. CRISPR-Cas9-guided knockout of the endogenous Jurkat CD4 co-receptor. a,** Flow cytometry shows Jurkat ROD6 cells (Cas9+ CD8+) after transfection with a gRNA targeting exon 5 in human CD4, resulting in 14% knockout efficiency. Cells in the CD4- gate were single-cell sorted and expanded in culture. **b,** Expanded clones (n = 19) were screened for expression of CD8α, CD4 and CD3 (not shown) via flow cytometry. **c,** PCR amplification of genomic DNA using a primer pair flanking human CD4 exon 5 (expected PCR product 660 bp) was performed on all clones. The PCR products of ROD6 cells and of six CD3$^{high}$ CD8+ CD4- clones (C10, D4, E4, E6, F3 and F4). were subjected to Sanger sequencing. Sequences derived from ROD6 cells aligned precisely with the reference human genome, while all selected clones (except clone C10) yielded forward and reverse chromatograms with unresolved peaks at the CD4 gRNA cut site, thus confirming targeted NHEJ. Clone E4 (renamed ROD10) was selected as the lead cell line moving forward.

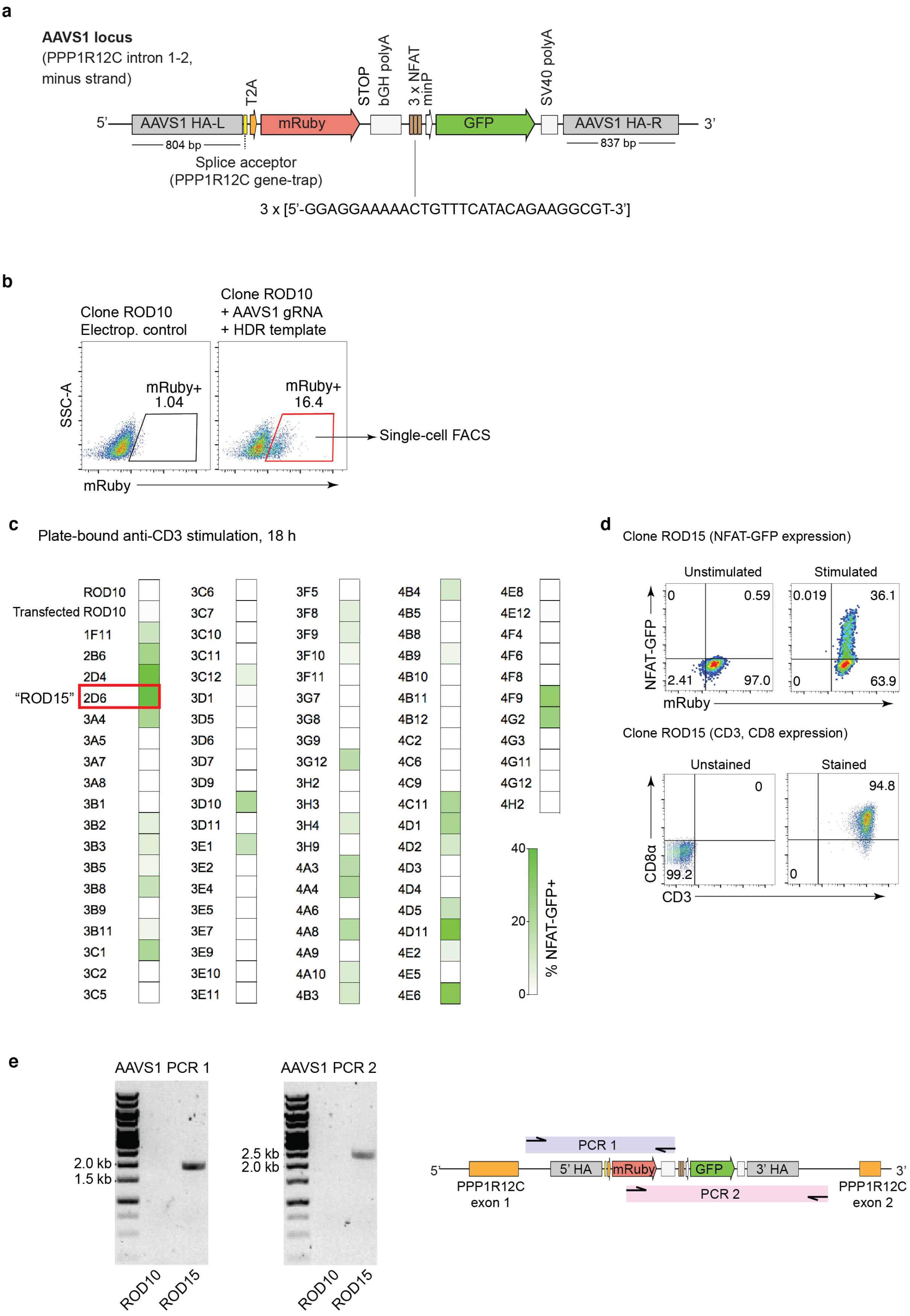
a
AAVS1 locus
(PPP1R12C intron 1-2,
minus strand)
T2A
STOP
bGH polyA
3 x NFAT
minP
SV40 polyA
5'
AAVS1 HA-L
mRuby
GFP
AAVS1 HA-R
3'
804 bp
837 bp
Splice acceptor
(PPP1R12C gene-trap)
3 x [5'-GGAGGAAAAACTGTTTCATACAGAAGGCGT-3']
b
Clone ROD10
Electrop. control
Clone ROD10
+ AAVS1 gRNA
+ HDR template
SSC-A
mRuby+
1.04
mRuby+
16.4
Single-cell FACS
mRuby
c
Plate-bound anti-CD3 stimulation, 18 h
ROD10
Transfected ROD10
1F11
2B6
2D4
"ROD15" 2D6
3A4
3A5
3A7
3A8
3B1
3B2
3B3
3B5
3B8
3B9
3B11
3C1
3C2
3C5
3C6
3C7
3C10
3C11
3C12
3D1
3D5
3D6
3D7
3D9
3D10
3D11
3E1
3E2
3E4
3E5
3E7
3E9
3E10
3E11
3F5
3F8
3F9
3F10
3F11
3G7
3G8
3G9
3G12
3H2
3H3
3H4
3H9
4A3
4A4
4A6
4A8
4A9
4A10
4B3
4B4
4B5
4B8
4B9
4B10
4B11
4B12
4C2
4C6
4C9
4C11
4D1
4D2
4D3
4D4
4D5
4D11
4E2
4E5
4E6
4E8
4E12
4F4
4F6
4F8
4F9
4G2
4G3
4G11
4G12
4H2
40
20
0
% NFAT-GFP+
d
Clone ROD15 (NFAT-GFP expression)
Unstimulated
Stimulated
NFAT-GFP
0
0.59
2.41
97.0
0.019
36.1
0
63.9
mRuby
Clone ROD15 (CD3, CD8 expression)
Unstained
Stained
CD8α
0
99.2
94.8
0
CD3
e
AAVS1 PCR 1
AAVS1 PCR 2
2.0 kb
1.5 kb
2.5 kb
2.0 kb
ROD10
ROD15
ROD10
ROD15
PCR 1
PCR 2
5'
5' HA
mRuby
GFP
3' HA
3'
PPP1R12C
exon 1
PPP1R12C
exon 2

**Supplementary Figure 4. CRISPR-Cas9 integration of an NFAT-GFP reporter of TCR signalling into the AAVS1 safe harbor locus. a,** Schematic representation of the designed HDR template utilized for the introduction of T2A-mRuby and NFAT-GFP genes into the AAVS1 locus. Expression of mRuby relied on the endogenous PPP1R12C promoter and correct splicing with PPP1R12C exon 1 (i.e., PPP1R12C gene-trap). The sequence of the tandem 3xNFAT:AP1 composite response element derived from the human IL-2 promoter is also shown[1]. **b,** Transfection of the ROD10 cell line (Cas9+ CD8+ CD4-) with AAVS1 gRNA and HDR template (PCR product) resulted in ~15% HDR efficiency, as measured by mRuby expression using flow cytometry. mRuby+ cells were single-cell sorted and expanded in culture. **c,** Heatmap shows NFAT-GFP expression levels (based on flow cytometry) from expanded clones (n = 84) that were stimulated overnight with plate-bound anti-human CD3e monoclonal antibody (OKT3). Clones expressing NFAT-GFP at > 10% frequencies were selected and screened for CD3, CD8 and GFP expression under resting conditions (not shown). **d,** The selected clone 2D6 (renamed ROD15) displays undetectable NFAT-GFP expression under resting conditions, and maintains high levels of CD3 and CD8 expression. **e,** Validation of targeted integration of T2A-mRuby and NFAT-GFP into the AAVS1 locus of ROD15 cells. PCR amplification of genomic DNA with primer pairs containing one primer outside the transfected HDR template revealed expected bands in ROD15 cells only. Sanger sequencing of PCR products 1 and 2 revealed correct sequences. PCR1 primers: RVL-136 and RVL-137, PCR 2 primers: RVL-139 and RVL-140.

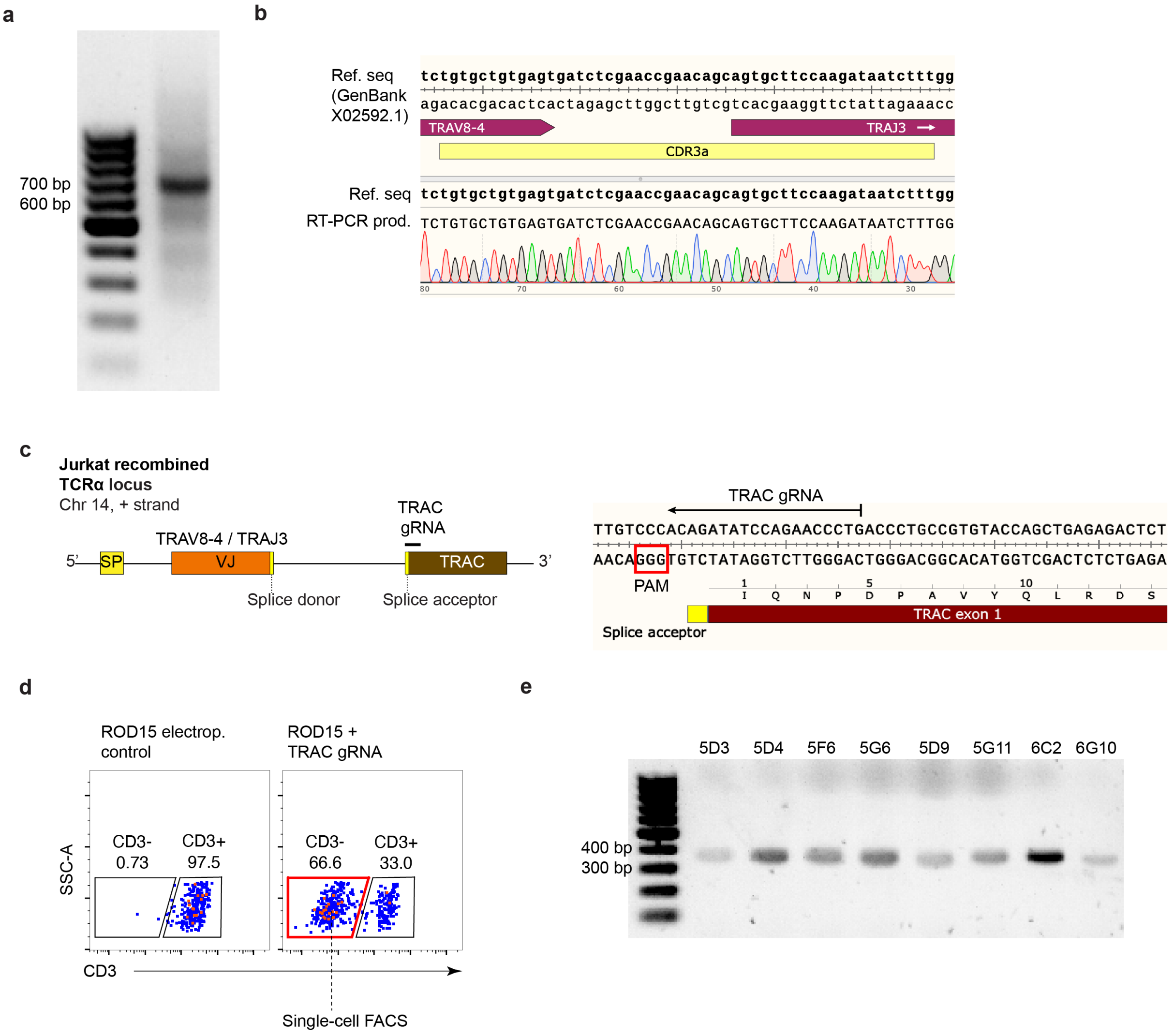


**f**

| | Clone | CD3+ (% of live) | CD3 MFI | NFAT-GFP+ (% of live) | Genotype | Outcome |
|---|---|---|---|---|---|---|
| "ROD20" | 5D3 | 0 | 81.3 | 56.8 | c.330_331insGCCCG | Premature stop codon |
| | 5D4 | 94 | 1220 | 42.8 | c.331_332insGGA | Two AA substitutions |
| | 5D9 | 43.9 | 287 | 27 | c.332_339del | Premature stop codon |
| | 5F6 | 0 | 72 | 26.9 | c.330_331insCCCAGGGG | Multiple AA substitutions |
| | 5G6 | 91.7 | 1065 | 44.8 | c.330_331insGGG | One AA substitution |
| | 5G11 | 60.9 | 482 | 46.9 | c.330_331insGATACG | Two AA insertions |
| | 6C2 | 93.9 | 1126 | 36.8 | c.T332G, c.T334A, c.C335A | Two AA substitutions |
| | 6G10 | 4.77 | 167 | 44 | c.331_339del | Premature stop codon |
| | ROD15 | 99.8 | 4862 | 40.6 | WT TRAC | N/A |

**Supplementary Figure 5. CRISPR-Cas9 knockout of the endogenous Jurkat TCRα chain. a,** PCR amplification of TSO-labelled Jurkat cDNA with ISPCR forward and TRAC reverse (RVL-70) primers. **b,** Gel extraction and Sanger sequencing of the ~ 700 bp band in **(a)** confirmed the expression of a single TCRα chain matching the Jurkat TCRα reference sequence (GenBank X02592.1). **c,** Schematic representation of the recombined Jurkat TCRα locus displaying the TRAC gRNA utilized for targeted knockout[2]. **d,** Flow cytometry shows that transfection of ROD15 cells (Cas9+ CD8+ CD4- mRuby+ NFAT-GFP) with TRAC gRNA results in ~ 65% TCRα knockout efficiency, as measured by surface expression of CD3. CD3 negative cells were single-cell sorted and expanded in culture. **e,** Expanded clones (n = 8) were subjected to RT-PCR using TRAV8-4 forward (RVL-69) and TRAC reverse (RVL-70) primers. All clones displayed a 350 bp band indicating that TRAC exon 1 skipping did not occur in any of them. **f,** Table summarizes Sanger sequencing results of RT-PCR products; 3-8 nt insertions or deletions at the TRAC gRNA target site was observed in most clones. Frameshifts leading to premature stop codons occurred in 3 out of 8 clones. Of these, clone 5D3 (renamed ROD20) had undetectable CD3 expression and robust NFAT-GFP expression in response to PMA-ionomycin, and was selected as the leading cell line moving forward. PAM: protospacer adjacent motif. TSO: template-switching oligonucleotide.

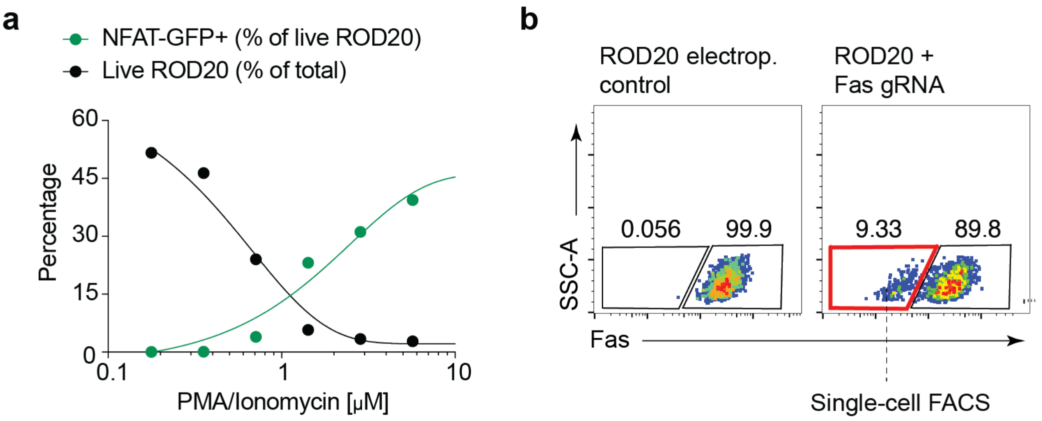


**c**

| | Clone | Fas+ (% of live) | CD3+ (% of live) | CD8+ (% of live) | % NFAT-GFP+ (stimulated) | % NFAT-GFP+ (resting) |
|---|---|---|---|---|---|---|
| "TnT" | 17B11 | 0.00 | 0.00 | 99.20 | 58.80 | 0.00 |
| | 17C1 | 0.00 | 0.00 | 99.90 | 79.30 | 0.00 |
| | 17D5 | 0.00 | 0.00 | 94.70 | 45.20 | 0.00 |
| | 17D6 | 0.00 | 0.00 | 99.50 | 89.20 | 0.00 |
| | 17E10 | 0.00 | 0.02 | 78.60 | 51.10 | 0.01 |
| | 17F4 | 0.00 | 0.00 | 84.70 | 1.69 | 0.00 |
| | 17F11 | 0.00 | 0.02 | 99.00 | 45.20 | 0.00 |
| | 17G2 | 0.00 | 0.00 | 96.80 | 80.30 | 0.00 |
| | 17G5 | 0.00 | 0.01 | 72.70 | 20.20 | 0.00 |
| | 17G8 | 0.00 | 0.00 | 97.70 | 54.10 | 0.00 |
| | 17H3 | 0.00 | 0.00 | 93.70 | 4.86 | 0.00 |
| | 17H7 | 0.00 | 0.00 | 84.90 | 75.10 | 0.00 |
| | ROD15 | 99.70 | 97.40 | 95.60 | ND | ND |

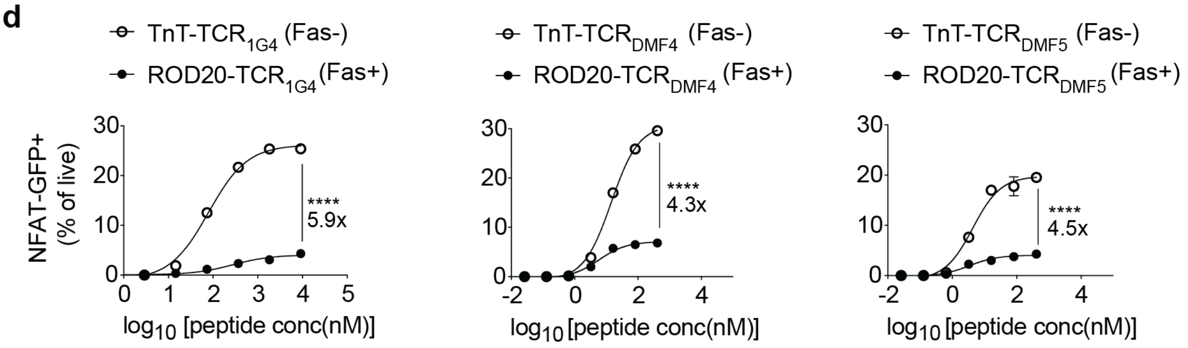


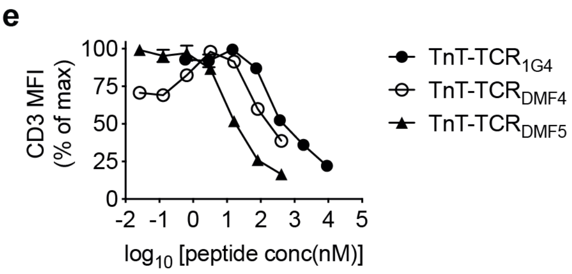

**Supplementary Figure 6. CRISPR-Cas9 knockout of Fas for the generation of TnT cells resistant to activation-induced cell death (AICD).** **a**, Increased cell death in the ROD20 cell line (Cas9+ CD8+ CD4- mRuby+ NFAT-GFP CD3-) after non-specific stimulation with increasing concentrations of PMA-ionomycin. This increase was inversely correlated to the proportion of NFAT-GFP-positive cells within surviving cells, thus indicating that increased cell death resulted from AICD. **b,** Transfection of ROD20 cells with Fas gRNA results in ~ 9% Fas knockout efficiency, as measured by flow cytometry. Fas negative cells were single-cell sorted and expanded in culture. **c,** Table summarizes phenotype of expanded clones (n = 12) based on flow cytometry measurement for expression of Fas, CD3, CD8α and NFAT-GFP (± PMA-iononomycin stimulation). Clone 17B11 was selected as the final version of the TnT platform. **d,** $TCR_{1G4}$, $TCR_{DMF4}$ and $TCR_{DMF5}$ were introduced into ROD20 (Fas+) or TnT (Fas-) cells via CRISPR-Cas9 HDR and co-cultured overnight with T2 cells pulsed with serially-diluted target peptides (n = 2). Activated TnT-TCR cells displayed significantly higher proportions of live NFAT-GFP-positive cells relative to ROD20-TCR cells, thus indicating increased resistance to AICD. **e**, CD3 mean fluorescence intensity (MFI) in TnT-TCR cells expressing $TCR_{1G4}$, $TCR_{DMF4}$ or $TCR_{DMF5}$ after co-culture T2 cells pulsed with serially-diluted target peptides (n = 2). Non-linear least squares fits are displayed in (**a**) and (**d**). Two-way ANOVA with Bonferroni post hoc test for multiple comparisons is displayed in (**d**), **** $P < 0.0001$.

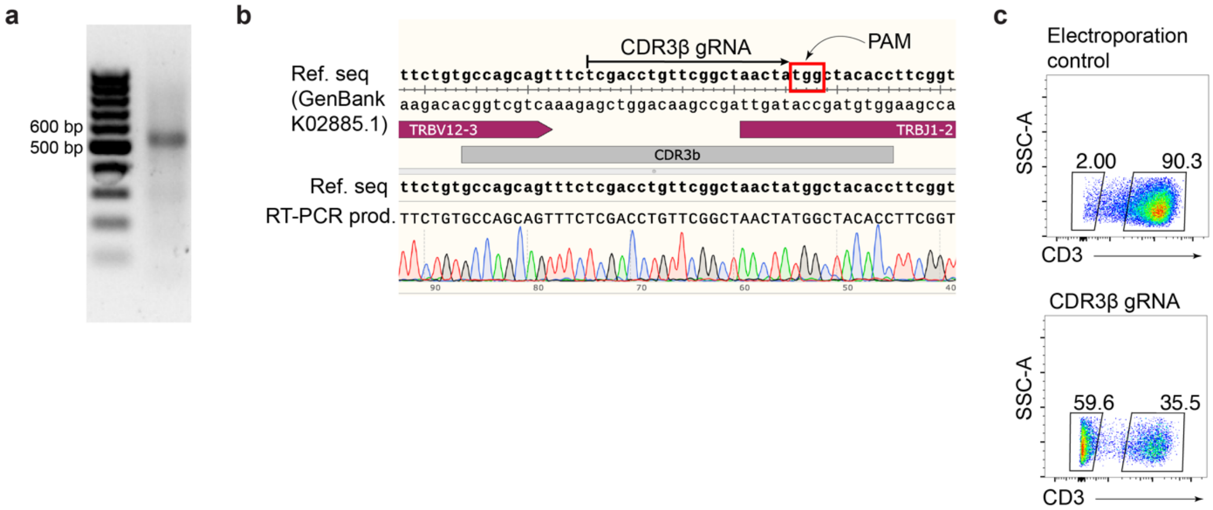


**Supplementary Figure 7. The recombined Jurkat CDRβ3 sequence in TCRβ is an ideal target for monoallelic CRISPR-Cas9 TCR reconstitution.** **a,** PCR amplification of TSO-labelled Jurkat cDNA with ISPCR forward and TRBC1/2 reverse (RVL-68) primers. **b,** Sanger sequencing of the ~ 550 bp band in **(a)** confirmed the expression of a single TCRβ chain matching the Jurkat TCRβ reference sequence (GenBank K02885.1). The designed CDR3β gRNA is also displayed. **c,** Flow cytometry shows transfection of Cas9+ Jurkat cells with CDR3β gRNA results in nearly 60% TCRβ knockout efficiency, as measured by surface expression of CD3. PAM: protospacer adjacent motif. TSO: template-switching oligonucleotide.

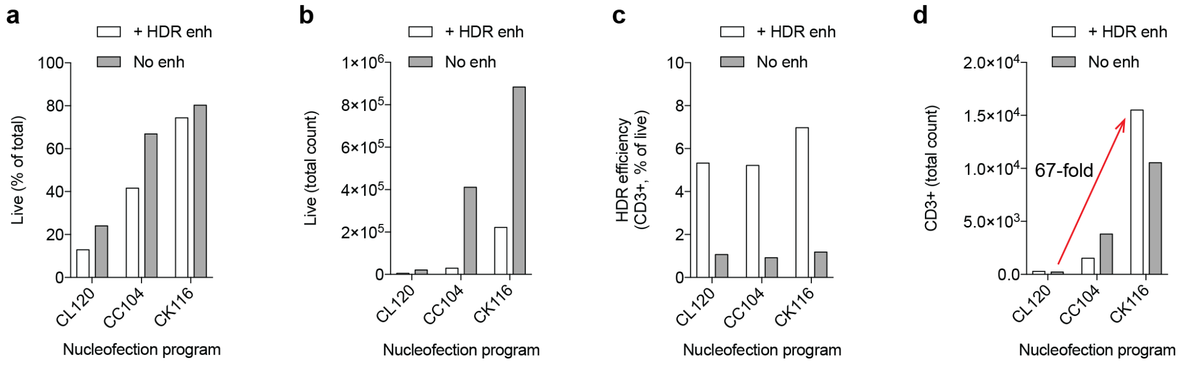


**Supplementary Figure 8. Optimization of homology-directed repair (HDR) efficiency for targeted TCR reconstitution in TnT cells.** 1 x $10^6$ TnT cells (CD3-) were transfected with 700 pmol of CDR3β gRNA and 1 μg of $TCR_{1G4}$ HDR template using three different electroporation programs: CL120, CC104 or CK116 (Lonza 4D nucleofector). Transfected samples were cultured with or without addition of HDR enhancer solution (30 μM final concentration, IDT) for the first 16 h following transfection. **a-b**, Cell viability (DAPI- cells) and **c-d**, HDR rates (CD3 restoration) were assessed by flow cytometry on day 5 post-transfection. **c**, Addition of HDR enhancer resulted in 5-7-fold increases in the percentages of CD3+ TnT cells across all nucleofection programs. **d**, Transfection of TnT cells with program CK116 in the presence of HDR enhancer led to a 67-fold increase in the total numbers of TCR-reconstituted cells relative to previously utilized conditions (CL120, no HDR enhancer).

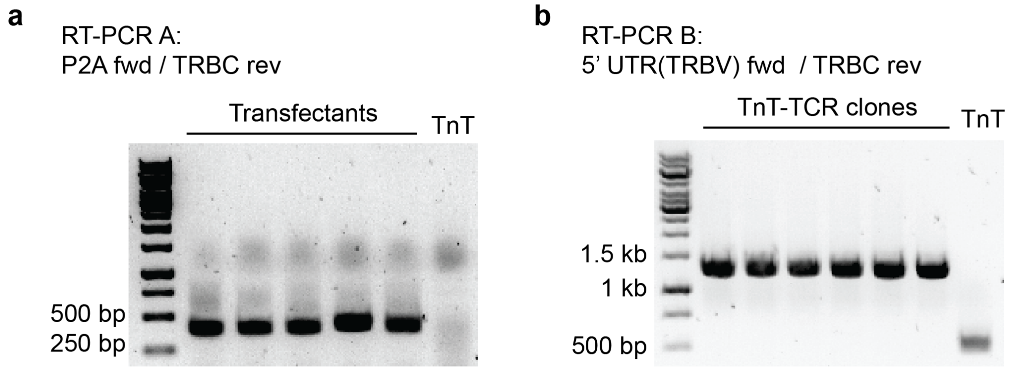


**Supplementary Figure 9. RT-PCR validation of CRISPR-Cas9 TCR reconstitution in TnT cells. a,** Gel shows results of RT-PCR using a forward primer annealing to the P2A sequence within the TCRαβ HDR template (RVL-144) and a reverse primer annealing to the endogenous Jurkat TRBC (RVL-145). Untransfected TnT cell samples display no PCR product, while samples derived from TCR transfectants display a 435 bp product consistent with targeted TCR integration and correct splicing with TRBC1. Products from RT-PCR were routinely utilized for deep sequencing of TCR selections (Genewiz Amplicon EZ). **b,** Gel shows results of RT-PCR using a forward primer annealing to the 5' UTR of the recombined Jurkat TCRβ VDJ exon (RVL-67c) and a reverse primer annealing to the endogenous Jurkat TRBC (RVL-68). Untransfected TnT cells show a 500 bp band corresponding to the endogenous Jurkat TCRβ, while TnT-TCR clones display a unique band at 1.3 kb demonstrating targeted integration of TCRαβ cassettes and their correct splicing with TRBC1 (see Fig.1b). Sanger sequencing of RT-PCR products was routinely performed for clone validation purposes.

**a**

**TCR alpha: TRAV21, TRAJ28**

```
A3       METLLGLLILWLQLQWVSSKQEVTQIPAALSVPEGENLVLNCSFTDSAIYNLQWFRQDPG  60
a3a      METLLGLLILWLQLQWVSSKQEVTQIPAALSVPEGENLVLNCSFTDSAIYNLQWFRQDPG  60
         ************************************************************
                     CDR2α
A3       KGLTSLLLIQSSQREQTSGRLNASLDKSSGRSTLYIAASQPGDSATYLCAVRPGGAGSYQ  120
a3a      KGLTSLLLVRPYQREQTSGRLNASLDKSSGRSTLYIAASQPGDSATYLCAVRPGGAGSYQ  120
         ********::  ************************************************

A3       LTFGKGTKLSVIP        133
a3a      LTFGKGTKLSVIP        133
         *************
```

**TCR beta: TRBV5-1, TRBJ2-7**

```
A3       MGSRLLCWVLLCLLGAGPVKAGVTQTPRYLIKTRGQQVTLSCSPISGHRSVSWYQQTPGQ  60
a3a      MGSRLLCWVLLCLLGAGPVKAGVTQTPRYLIKTRGQQVTLSCSPISGHRSVSWYQQTPGQ  60
         ************************************************************

A3       GLQFLFEYFSETQRNKGNFPGRFSGRQFSNSRSEMNVSTLELGDSALYLCASSPNMADEQ  120
a3a      GLQFLFEYFSETQRNKGNFPGRFSGRQFSNSRSEMNVSTLELGDSALYLCASSPNMADEQ  120
         ************************************************************

A3       YFGPGTRLTVT  131
a3a      YFGPGTRLTVT  131
         ***********
```

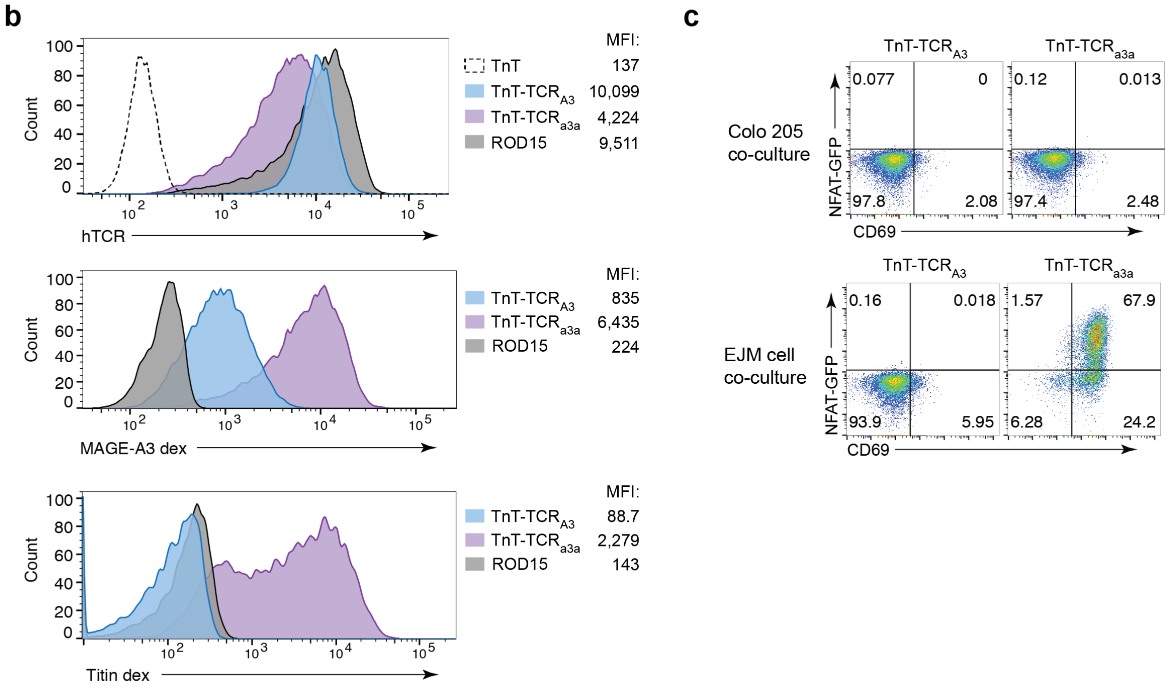


**Supplementary Figure 10. Characterization of TnT cells with TCRs recognizing the MAGE-A3 peptide antigen. a,** Protein sequence alignments of TCR$_{A3}$ and its phage display-engineered variant TCR$_{a3a}$[3-5]. The four substitutions introduced into the CDR2α of TCR$_{a3a}$ are highlighted in red[6]. **b,** TnT clones expressing TCR$_{A3}$ and TCR$_{a3a}$ were generated by CRISPR-Cas9 HDR followed by single-cell FACS. Levels of TCR expression (anti-TCR monoclonal antibody clone IP26), MAGE-A3 peptide-MHC dextramer binding and titin peptide-MHC dextramer binding in TnT-TCR$_{A3}$, TnT-TCR$_{a3a}$ and negative control cell lines are displayed. **c,** Flow cytometry of NFAT-GFP and CD69 expression after overnight co-culture of TnT-TCR$_{A3}$ and TnT-TCR$_{a3a}$ cells with Colo 205 (HLA-A*0101 MAGE-A3-) and EJM (HLA-A*0101+ MAGE-A3+) cancer cell lines.

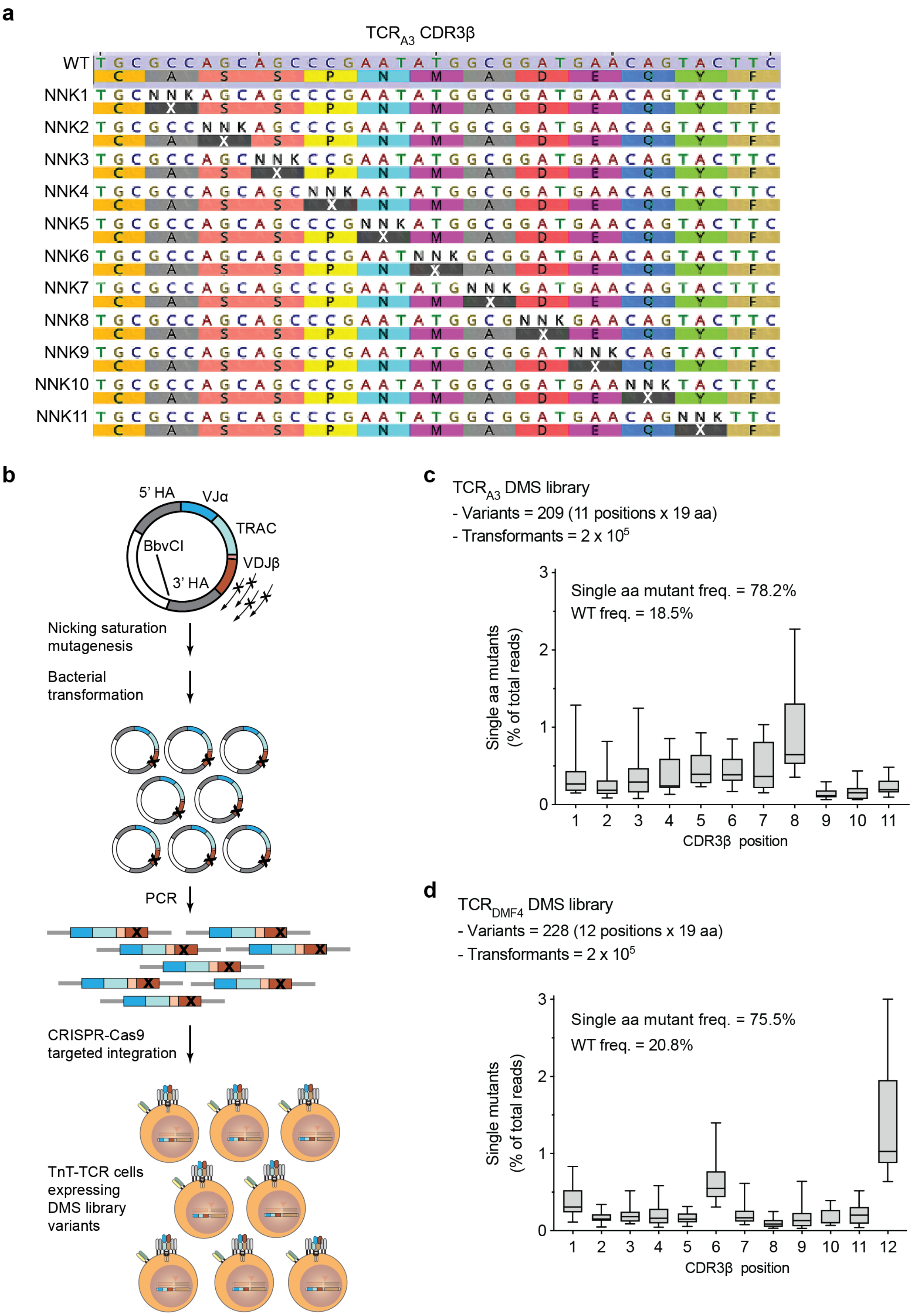
a
TCR$_{A3}$ CDR3β
WT
NNK1
NNK2
NNK3
NNK4
NNK5
NNK6
NNK7
NNK8
NNK9
NNK10
NNK11
b
5' HA
VJα
TRAC
BbvCI
VDJβ
3' HA
Nicking saturation mutagenesis
Bacterial transformation
PCR
CRISPR-Cas9 targeted integration
TnT-TCR cells expressing DMS library variants
c
TCR$_{A3}$ DMS library
- Variants = 209 (11 positions x 19 aa)
- Transformants = 2 x 10$^5$
Single aa mutant freq. = 78.2%
WT freq. = 18.5%
Single aa mutants (% of total reads)
CDR3β position
d
TCR$_{DMF4}$ DMS library
- Variants = 228 (12 positions x 19 aa)
- Transformants = 2 x 10$^5$
Single aa mutant freq. = 75.5%
WT freq. = 20.8%
Single mutants (% of total reads)
CDR3β position

**Supplementary Figure 11. Generation of CDR3β deep mutational scanning (DMS) libraries by plasmid-based nicking saturation mutagenesis. a,** The CDR3β of $TCR_{A3}$ (displayed) and $TCR_{DMF4}$ were targeted for DMSbyintroducing NNK degenerate codons tiled across each position. **b,** Plasmids containing TCRαβ cassettes encoding $TCR_{A3}$ or $TCR_{DMF4}$ were subjected to saturation mutagenesis using the method from Wrenbeck et al. 2016[7]. TnT-TCR cells expressing unique TCR variants were isolated by FACS based on restoration of CD3 surface expression. **c-d,** PCR amplicons generated from $TCR_{A3}$ and $TCR_{DMF4}$ plasmid DMS libraries were subjected to deep sequencing for library validation. The frequency of single amino acid mutants according to their position in CDR3β, the frequency of all single amino acid mutants and the frequency of wild-type TCR in each library are displayed. In line with reported method performance, both $TCR_{A3}$ and $TCR_{DMF4}$ DMS libraries contained every possible single amino acid substitution ($TCR_{A3}$ = 209 variants, $TCR_{DMF4}$ = 228 variants), with single mutants accounting for 75.5% to 78.2% of sequencing reads and background parental TCR sequences accounting for 18.5% to 20.8% of sequencing reads.

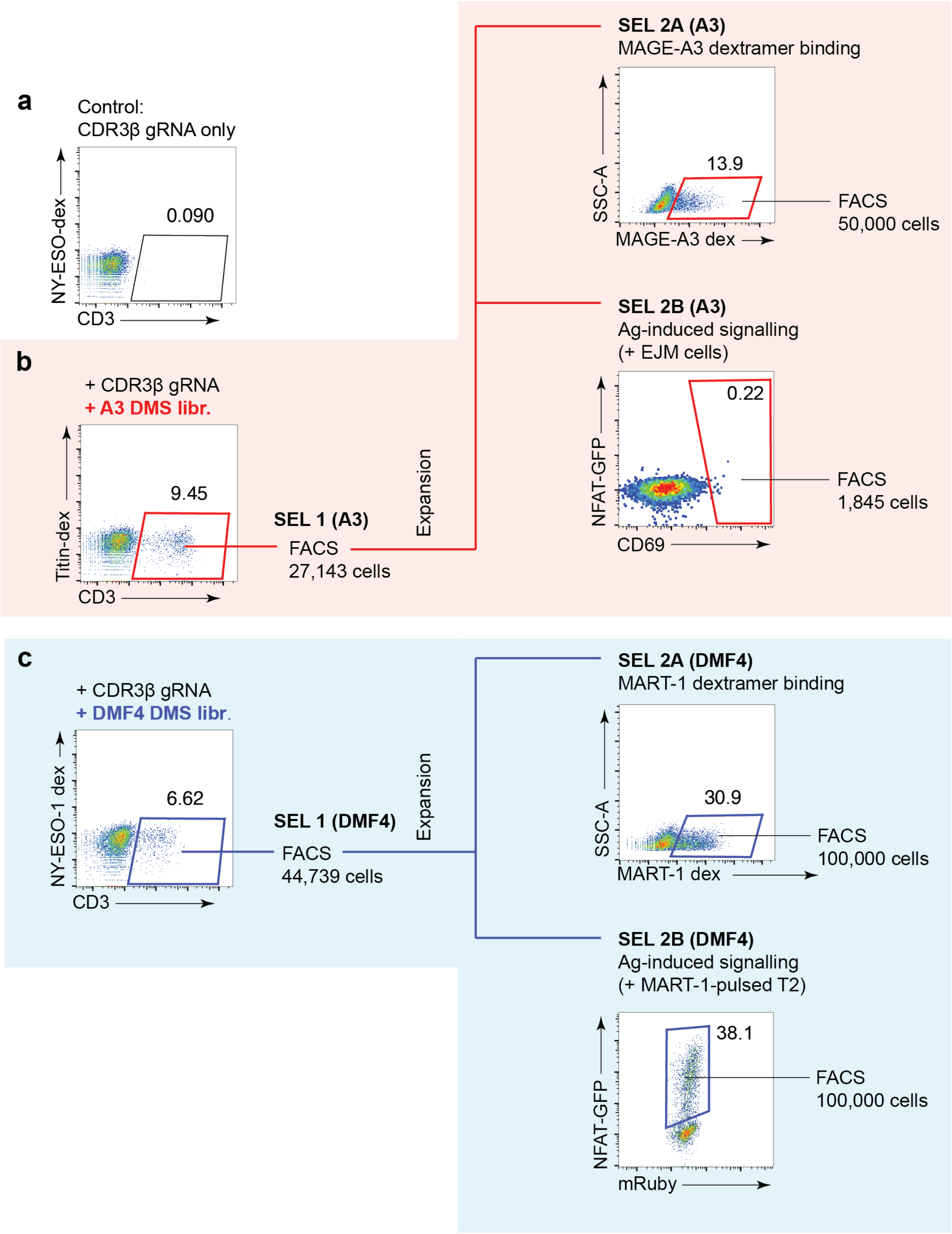


**Supplementary Figure 12. Binding- and signaling-based selection strategies for screening TCR$_{A3}$ and TCR$_{DMF4}$ DMS libraries. a,** TnT cells transfected with CDR3β gRNA display no surface expression of CD3 and no binding to a control NY-ESO-1 pMHC dextramer (HLA-A*0201, SLLMWITQC). **b,** TnT cells were transfected with CDR3β gRNA and HDR templates encoding the TCR$_{A3}$ DMS library. FACS selections based on restored CD3 surface expression (SEL 1), binding to MAGE-A3 pMHC dextramer (SEL 2A) and activation following co-culture with MAGE-A3+ EJM cells (SEL 2B) are displayed. **c**, TnT cells were transfected with CDR3β gRNA and HDR templates encoding the TCR$_{DMF4}$ DMS library. FACS selections based on restored CD3 surface expression (SEL 1), binding to ELAGIGILTV pMHC dextramer (SEL 2A) and activation following co-culture with ELAGIGILTV-pulsed T2 cells (SEL 2B) are displayed. In addition to anti-CD3 antibody, transfected cells were co-stained with HLA-matched control peptide-MHC dextramers but no cross-reactive TnT-TCR cells were detected.

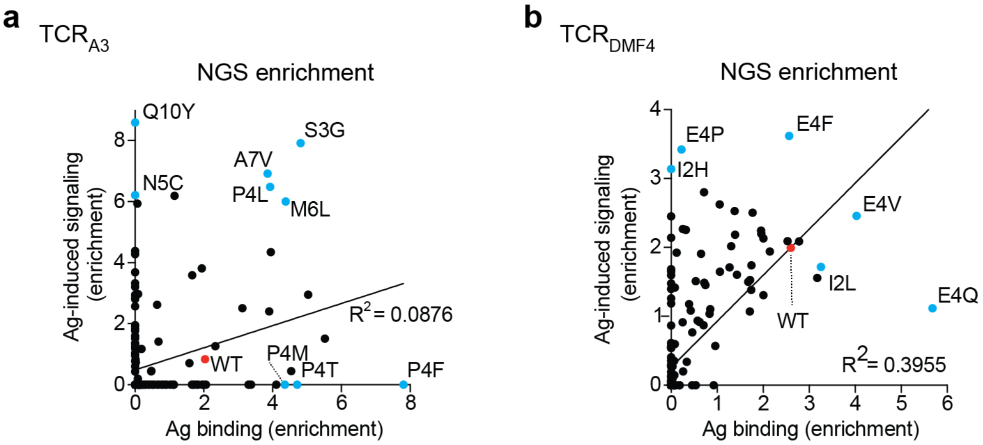


**Supplementary Figure 13. Differential enrichment of TCR CDR3β single mutants in binding-based and signaling-based selections. a**, **b**, Graphs display enrichment based on deep sequencing reads of individual $TCR_{A3}$ (**a**) or $TCR_{DMF4}$ (**b**) variants in DMS SEL 2A (peptide-MHC binding) and SEL 2B (antigen-induced signalling). Variants selected for further validation are highlighted in blue (see Fig. 2e-i), wild-type $TCR_{A3}$ and $TCR_{DMF4}$ are highlighted in red. Linear regression analysis is displayed.

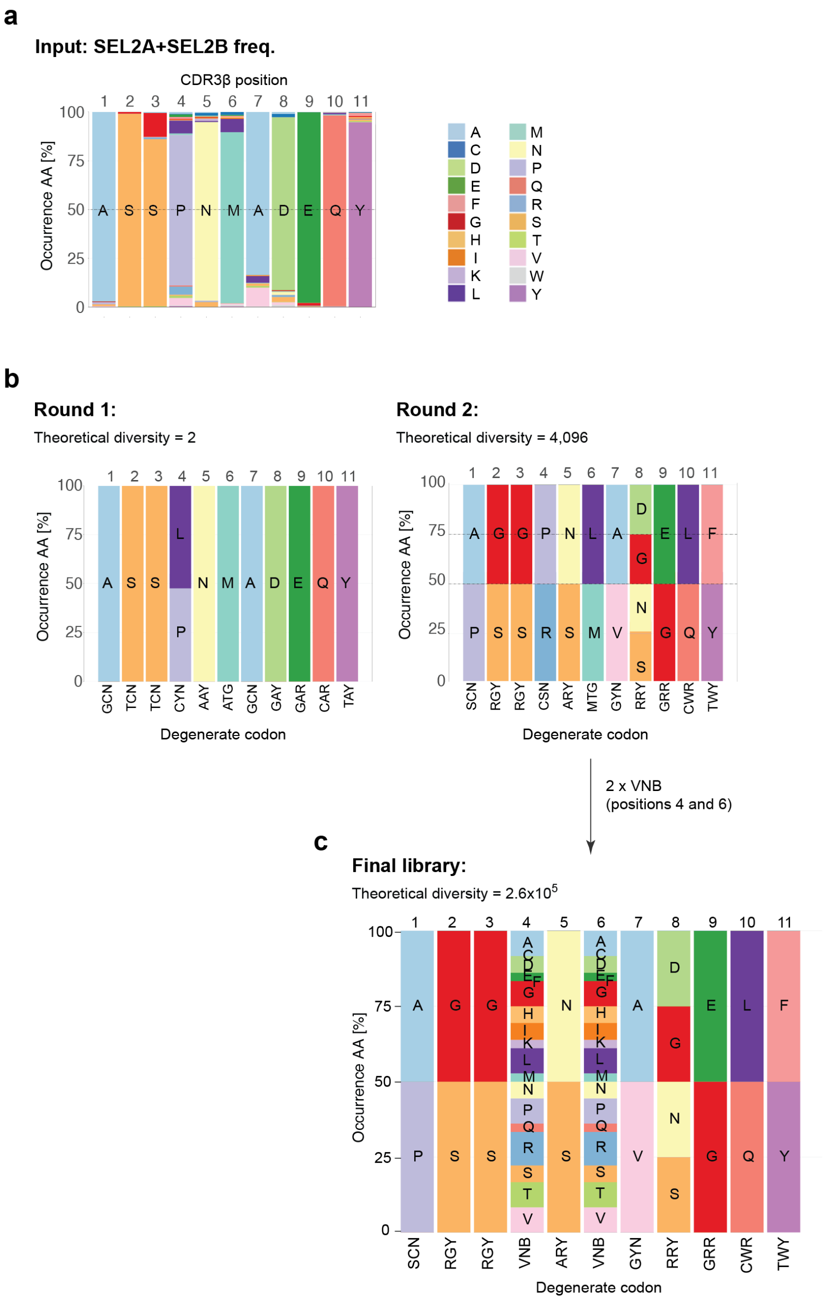


**Figure S14. Combinatorial library design based on DMS of the TCRA3 CDR3β. a,** Bar plots show the frequencies of single mutants present in TCRA3 DMS selections 2A (MAGE-A3 peptide-MHC+) and 2B (NFAT-GFP+) (added and normalized to 100%). **b,** Pooled DMS data was used as input for our previously described algorithm[8] with the aim of re-capitulating observed amino acid frequencies with degenerate codons (two iterations with varying sensitivity to lower frequency amino acids are displayed). **c,** To take advantage of the full attainable diversity of mammalian cell display, the "Round 2" solution (theoretical diversity of 4,096 sequences) was rationally modified to include two "VNB" codons at positions 4 and 6 (high levels of enrichment in DMS), leading to a theoretical diversity of 2.6 x $10^5$ variants. Degenerate base symbols: R = A, G; Y = C, T; S = G, C; W = A, T; K = G, T; M = A, C; B = C, G, T; D = A, G, T; H = A, C, T; V = A, C, G; N = any base.

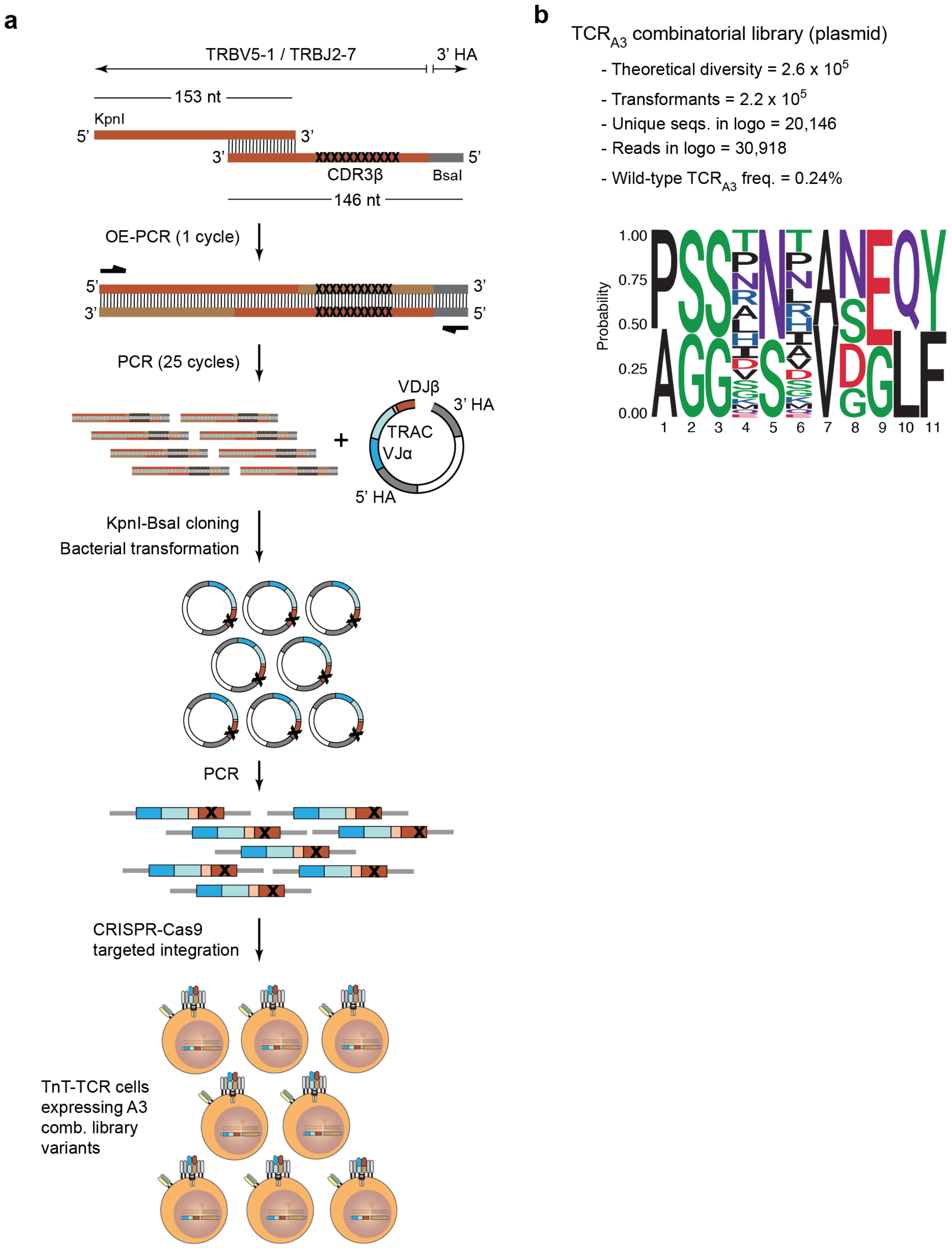
a
TRBV5-1 / TRBJ2-7
3' HA
153 nt
KpnI
5'
3'
3'
5'
CDR3β
BsaI
146 nt
OE-PCR (1 cycle)
5'
3'
3'
5'
PCR (25 cycles)
VDJβ
3' HA
TRAC
VJα
5' HA
+
KpnI-BsaI cloning
Bacterial transformation
PCR
CRISPR-Cas9
targeted integration
TnT-TCR cells
expressing A3
comb. library
variants
b
TCRA3 combinatorial library (plasmid)
- Theoretical diversity = 2.6 x 10^5
- Transformants = 2.2 x 10^5
- Unique seqs. in logo = 20,146
- Reads in logo = 30,918
- Wild-type TCRA3 freq. = 0.24%
Probability
1.00
0.75
0.50
0.25
0.00
1 2 3 4 5 6 7 8 9 10 11

**Supplementary Figure 15. Generation, characterization and selection of TCR$_{A3}$ combinatorial libraries. a,** Overlapping single-stranded oligodeoxynucleotides (ssODN) containing KpnI (TRBV5-1) or BsaI (3' homology arm) restriction sites were designed to build the TCR$_{A3}$ combinatorial library. Reverse ssODNs contained minus-strand degenerate codons complementary to the original library design. Double-stranded DNA was generated by overlap-extension PCR and flanking primers were utilized to further PCR-amplify the resulting product. Libraries were assembled by restriction cloning into a pJurTCRb plasmid containing the TCR$_{A3}$ TCRαβ cassette. After transformation into bacteria, plasmids were purified and amplified by PCR in order to generate dsDNA HDR templates for transfection into TnT cells. TnT-TCR cells expressing unique TCR variants were isolated by FACS based on restoration of CD3 surface expression. **b**, PCR amplicons generated from plasmid preparations were subjected to deep sequencing for library validation. The frequencies of specific amino acids at each CDR3β position are displayed (logo weighted on unique variant frequency).

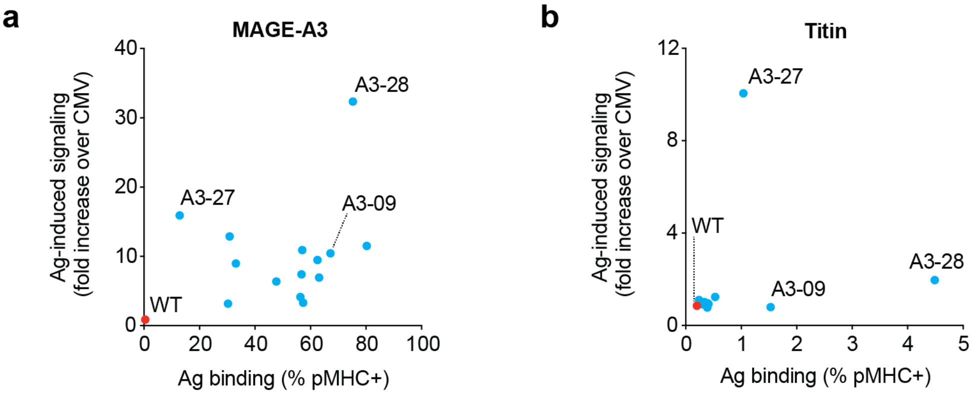


**Supplementary Figure 16. Differential antigen binding and antigen-induced signaling in engineered TCR$_{A3}$ variants.** TnT cells expressing wild-type TCR$_{A3}$ and selected TCR$_{A3}$ combinatorial variants were assessed for: (1) binding to MAGE-A3 peptide-MHC and titin peptide-MHC dextramers, and (2) for activation after co-culture with Colo 205 cells pulsed with MAGE-A3 or titin peptides (fold increase in the proportion of CD69$^{high}$ TnT-TCR cells). Graphs display the correlation between antigen-binding and antigen-induced signaling in TCR$_{A3}$ combinatorial variants in the context of MAGE-A3 (**a**) and titin (**b**) epitopes. TCR$_{A3}$ variants displaying cross-reactivity to titin are highlighted in blue. Graph generated from data shown in Figs. 3h, 3i and 4e.

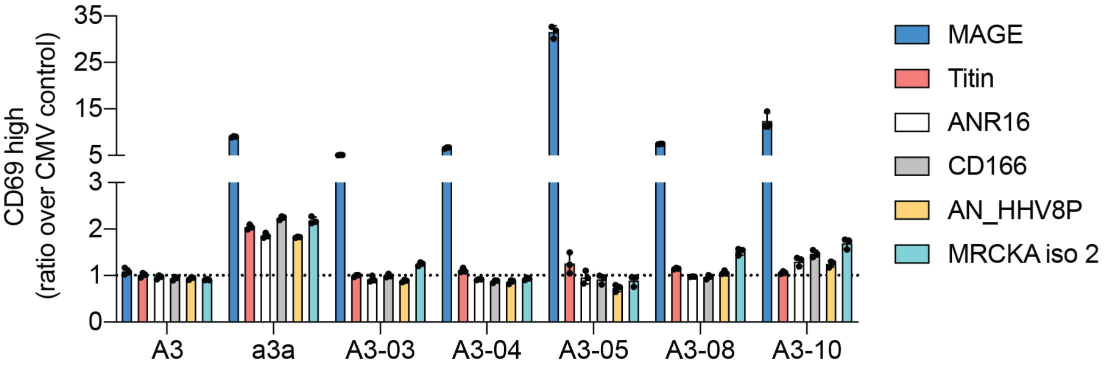


**Supplementary Figure 17. Cross-reactivity of selected TnT-$TCR_{A3}$ clonal variants against Colo 205 cells pulsed with predicted $TCR_{a3a}$ off-targets.** TnT-TCR clones expressing $TCR_{A3}$, $TCR_{a3a}$, $TCR_{A3\text{-}03}$, $TCR_{A3\text{-}04}$, $TCR_{A3\text{-}05}$, $TCR_{A3\text{-}08}$ or $TCR_{A3\text{-}10}$ were generated by CRISPR-Cas9 HDR followed by single-cell FACS of CD3+ cells. Expanded clones (4-6 per TCR) were screened by RT-PCR and flow cytometry in order to validate correct transgenic TCR sequence; maintained CD3, CD8 and NFAT-GFP (post-stimulation) expression; and ability to bind MAGE-A3 pMHC dextramer. Selected TnT-TCR clones were co-cultured with Colo 205 cells pulsed with predicted $TCR_{a3a}$ off-target peptides or with negative control CMV peptide . After overnight co-culture, the percentage of $CD69^{high}$ cells for each TnT-TCR cell line was determined by flow cytometry and normalized to its respective background. Data are displayed as mean ±SD, n = 3.

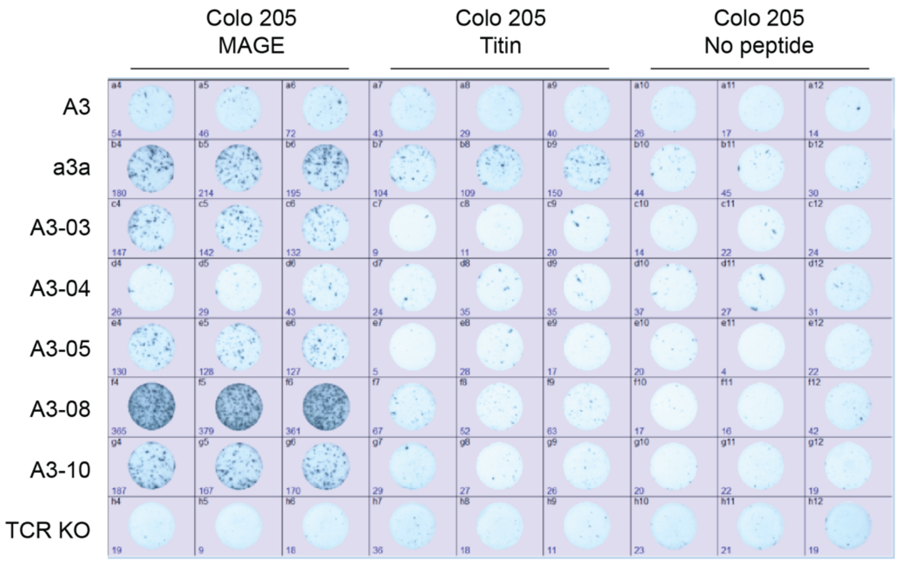


**Supplementary Figure 18. Selected $TCR_{A3}$ variants display negligible titin-induced activation.** ELISpot images corresponding to data in figure 5-f.

**a**

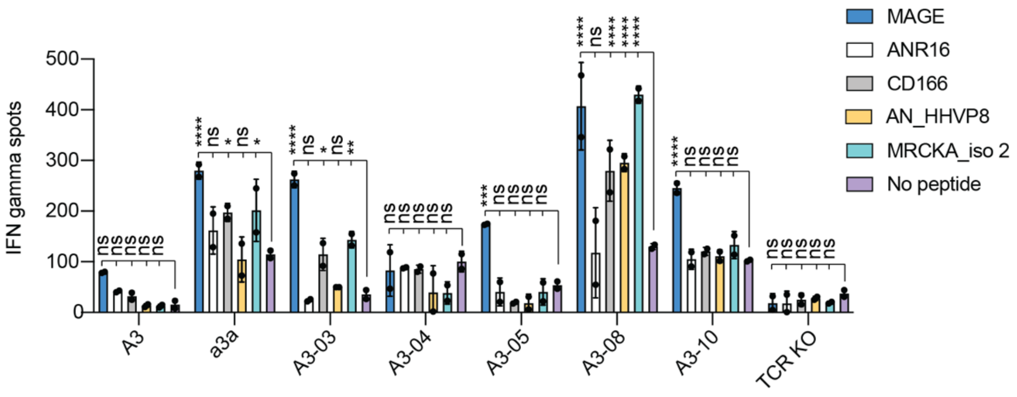


**b**

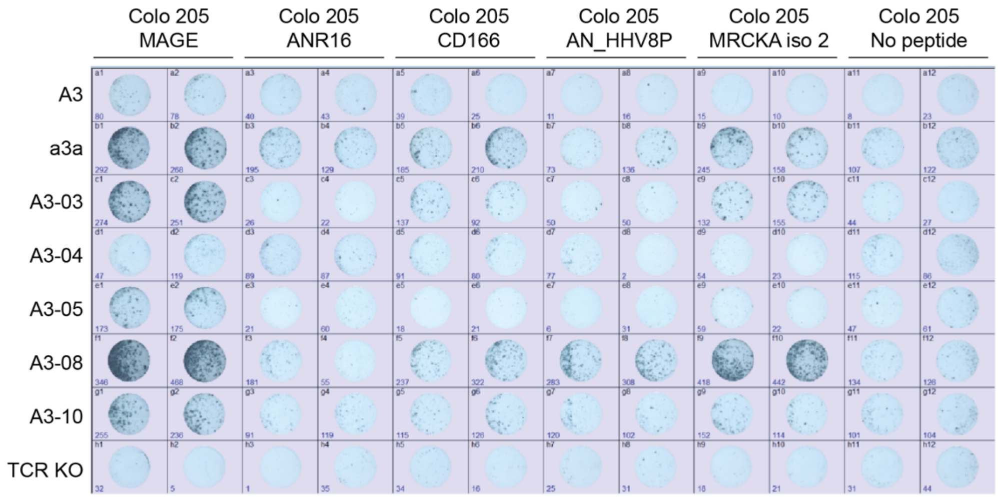


**Supplementary Figure 19. Primary T cells expressing TCR$_{A3-05}$ and TCR$_{A3-10}$ TCRs are highly selective for the MAGE-A3$_{168-176}$ EVDPIGHLY peptide.** Primary CD8+ human T cells were transfected with Cas9 RNP complexes targeting the TRAC and TRBC regions, alongside one of the following TCR-encoding HDR templates: TCR$_{A3}$, TCR$_{a3a}$, TCR$_{A3-03}$, TCR$_{A3-04}$, TCR$_{A3-05}$, TCR$_{A3-08}$ or TCR$_{A3-10}$. A no-HDR-template control (i.e., TCR KO) was also included. Transfectants were co-cultured overnight with peptide-pulsed Colo 205 cells (n = 2, 2.5x10$^{5}$ T cells per well). **a,** Activation of T cells expressing transgenic TCRs as assessed by IFN-γ ELISpot. **b,** Image corresponding to data in (**a**). Two-way ANOVA with Bonferroni post hoc test for multiple comparisons against non-peptide-pulsed controls is displayed; * $P$ < 0.05, ** $P$ < 0.01, *** $P$ < 0.001, **** $P$ < 0.0001, ns = not significant. RNP: ribonucleoprotein.

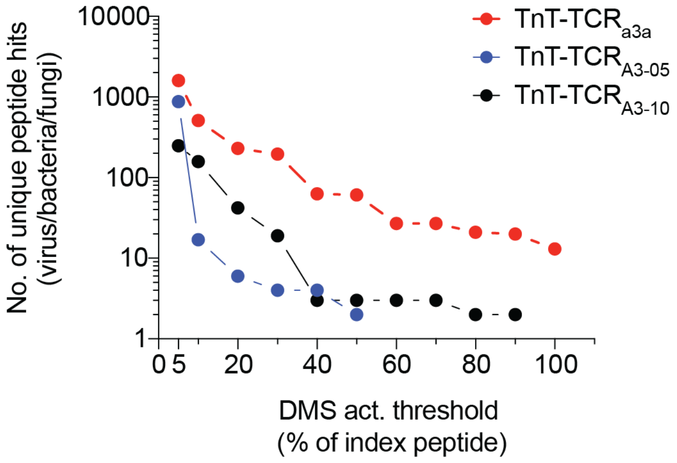


**Supplementary Figure 20. TCR$_{A3\text{-}05}$ and TCR$_{A3\text{-}10}$ have a reduced number of predicted non-human peptide hits.** TnT-TCR$_{a3a}$, TnT-TCR$_{A3\text{-}05}$ and TnT-TCR$_{A3\text{-}10}$ cells were co-cultured overnight with Colo 205 cells pulsed with single mutants of the MAGE-A3 wild-type peptide (i.e., peptide DMS library, n = 171). The sequences of peptides mediating 5, 10, 20, 30, 40, 50, 60, 70, 80, 90 and 100 percent activation relative to the wild-type peptide were utilized to generate motifs to query the UniProtKB database. Displayed are the number of unique bacterial, viral and fungal peptide hits for each TCR. TnT-TCR$_{A3\text{-}05}$ motifs above the 50% activation threshold returned no hits. TnT-TCR$_{A3\text{-}10}$ motifs above the 90% activation threshold returned no hits.

### Supplementary Tables 1-7

**Supplementary Table 1.** Previously discovered TCRs and target peptides utilized in this study

| Name | Target protein(s) | Target peptide | MHC restriction | Peptide sequence | Comments |
|---|---|---|---|---|---|
| TCR$_{1G4}$ | NY-ESO-1 | NY-ESO-1$_{157-164}$ | HLA-A*0201 | SLLMWITQC | |
| TCR$_{DMF4}$ | MART-1 | MART-1$_{26-35(27L)}$ | HLA-A*0201 | ELAGIGILTV | Anchor-modified peptide |
| TCR$_{DMF5}$ | MART-1 | MART-1$_{26-35(27L)}$ | HLA-A*0201 | ELAGIGILTV | Anchor-modified peptide |
| TCR$_{A3}$ | MAGE-A3 | MAGE-A3$_{168-176}$ | HLA-A*0101 | EVDPIGHLY | |
| TCR$_{a3a}$ | MAGE-A3 | MAGE-A3$_{168-176}$ | HLA-A*0101 | EVDPIGHLY | |
| | Titin | Titin$_{24,337-24,345}$ | HLA-A*0101 | ESDPIVAQY | |

**Supplementary table 2.** Unique clones identified in SEL 3A (MAGE-A3-induced signalling) and SEL 3B (titin-induced signalling) by means of deep sequencing

| Total clones SEL 3A + 3B | Clones in SEL 3A only | Clones in SEL 3B only | Clones in both SEL 3A and SEL 3B | Clones enriched >2-fold in SEL 3A and not enriched in SEL 3B (enrich. over SEL 1) |
|---|---|---|---|---|
| 1600 | 910 | 365 | 325 | 195 |

**Supplementary table 3.** Top-ranked $TCR_{A3}$ variants with predicted high specificity for MAGE-A3. Experimentally validated variants are highlighted in bold.

| TCR | SEL 2 (MAGE) enrichment over SEL 1 | SEL 3A (MAGE) enrichment over SEL 1 | SEL 3B (titin) enrichment over SEL 1 | SEL 3A frequency rank | SEL 3A enrichment rank | Score |
|---|---|---|---|---|---|---|
| **A3-001** | **41.9** | **130.5** | **0.0** | **45** | **8** | **53** |
| **A3-002** | **54.8** | **103.1** | **0.0** | **59** | **13** | **72** |
| **A3-003** | **53.1** | **78.2** | **0.0** | **76** | **19** | **95** |
| **A3-004** | **82.1** | **78.1** | **0.0** | **77** | **20** | **97** |
| **A3-005** | **25.9** | **65.7** | **0.0** | **21** | **24** | **45** |
| **A3-006** | **71.7** | **65.6** | **0.0** | **89** | **25** | **114** |
| **A3-007** | **43.5** | **63.3** | **0.0** | **93** | **26** | **119** |
| **A3-008** | **35.4** | **57.0** | **0.0** | **97** | **31** | **128** |
| **A3-009** | **32.2** | **55.9** | **0.0** | **99** | **33** | **132** |
| **A3-010** | **88.6** | **53.5** | **0.0** | **102** | **34** | **136** |
| A3-011 | 19.1 | 48.9 | 0.0 | 31 | 40 | 71 |
| **A3-012** | **25.0** | **46.0** | **0.4** | **8** | **45** | **53** |
| A3-013 | 39.5 | 43.4 | 0.0 | 124 | 50 | 174 |
| A3-014 | 23.4 | 42.2 | 0.0 | 127 | 53 | 180 |
| A3-015 | 20.9 | 41.0 | 0.0 | 132 | 55 | 187 |
| A3-016 | 33.8 | 41.0 | 0.0 | 133 | 56 | 189 |
| A3-017 | 26.1 | 40.4 | 0.0 | 70 | 58 | 128 |
| A3-018 | 35.6 | 39.1 | 0.0 | 40 | 60 | 100 |
| A3-019 | 29.0 | 38.5 | 0.0 | 136 | 61 | 197 |
| A3-020 | 25.8 | 37.3 | 0.0 | 139 | 63 | 202 |
| A3-021 | 26.1 | 34.7 | 0.0 | 83 | 68 | 151 |
| A3-022 | 24.5 | 33.5 | 0.0 | 43 | 74 | 117 |
| A3-023 | 9.2 | 31.6 | 0.0 | 92 | 76 | 168 |
| A3-024 | 11.4 | 22.1 | 0.0 | 41 | 98 | 139 |
| **A3-025** | **12.1** | **21.8** | **0.5** | **88** | **99** | **187** |
| A3-026 | 15.8 | 19.3 | 0.0 | 62 | 110 | 172 |
| **A3-027** | **4.9** | **17.9** | **0.2** | **50** | **117** | **167** |
| **A3-028** | **13.6** | **15.4** | **1.0** | **11** | **133** | **144** |
| A3-029 | 10.3 | 15.4 | 0.1 | 29 | 134 | 163 |

**Supplementary table 4.** Motifs generated from discrete activation thresholds following DMS of MAGE-A3$_{168\text{-}176}$ EVDPIGHLY target peptide

| TCR | Thresh. % | Motif |
|---|---|---|
| a3a | 5 | E-[ACFGHILMNQSTV]-[DY]-[FPW]-[ILMV]-[ACDEFGHILMNPQRSTVWY]-[ACDEFGHIKLMNPQRSTVWY]-[ACDEFGHIKLMNPQRSTVWY]-[CKLMTVY] |
| | 10 | E-[ACFGHILMNQSTV]-[DY]-[PW]-[ILMV]-[ACDEFGHILMNPQRSTVWY]-[ACDEFGHIKLMNPQRSTVWY]-[ACDEFGHIKLMNPQRSTVWY]-[MTVY] |
| | 20 | E-[ACGILMNQSTV]-[DY]-[PW]-[ILMV]-[ACDEFGHILMNPQSTVWY]-[ACEFGHIKLMNPQSTVWY]-[ACDEFGHIKLMNPQRSTVWY]-[VY] |
| | 30 | E-[ACGILMQSTV]-[DY]-[PW]-[ILMV]-[ACDEFGHILMNPQSTVWY]-[ACEFGHILMNPQSTVWY]-[ACDEFGHIKLMNPQRSTVWY]-[VY] |
| | 40 | E-[ACGILMQSTV]-[DY]-P-[ILMV]-[ACDEFGHILMNPQSTVWY]-[ACEFGHILMNPQSTVWY]-[ACDEFGHIKLMNPQRSTVWY]-Y |
| | 50 | E-[ACGILMQSTV]-[DY]-P-[ILMV]-[ACDEFGHILMNPQSTVY]-[ACEFGHILMNQSTVWY]-[ACDEFGHIKLMNPQRSTVWY]-Y |
| | 60 | E-[AILMQSTV]-D-P-[ILM]-[ACDEFGHILMNPQSTVY]-[AEFGHILMNQSTVWY]-[ACDEFGHIKLMNPQRSTVWY]-Y |
| | 70 | E-[AILMQSTV]-D-P-[ILM]-[ACDEFGHILMNPQSTVY]-[AEFGHILMNQSTVWY]-[ACDEFGHIKLMNPQRSTVWY]-Y |
| | 80 | E-[AILMSTV]-D-P-[ILM]-[ADEFGHILMNPQSTVY]-[AEFHILMQSTVW]-[ACDEFHKLMNPQRSTWY]-Y |
| | 90 | E-[AILMSTV]-D-P-[ILM]-[DEFGHILMNPSTVY]-[AEFHILMQSTVW]-[ACDEFHLMNPRSTY]-Y |
| | 100 | E-[AILMSTV]-D-P-[ILM]-[DEFGHILMNSTY]-[AEFHILQSTVW]-[ACDEFHLMNPRSY]-Y |
| A3-005 | 5 | [CDEFGHILNPQRS]-[ACFGILMNQSTV]-[DY]-P-[IMV]-[ACDEFGHILMNQSTY]-[ACEFGHILMNPQSTVW]-[ACDEFGHIKLMNPQRSTVWY]-[VY] |
| | 10 | E-[ACGILMNQSTV]-[DY]-P-[IM]-[CDEFGHILMNQST]-[AEFHILNPQSTVW]-[ACDEFGHILMNPQRSTVWY]-Y |
| | 20 | E-[ACGILMQSTV]-[DY]-P-[IM]-[DEGILMNST]-[AEHILSVW]-[ACDEFGHILMNPQRSTVWY]-Y |
| | 30 | E-[ACILMSTV]-D-P-[IM]-[DEGMNT]-[AEHSVW]-[ACDEFHILMNPQSTWY]-Y |
| | 40 | E-[ACILMSTV]-D-P-[IM]-[DEGNT]-[AEHW]-[ACDEFHLMNQSWY]-Y |
| | 50 | E-[AILMSTV]-D-P-[IM]-[DEGNT]-[AEHW]-[CDEFHLMNSWY]-Y |
| | 60 | E-[AILMSTV]-D-P-[IM]-[DEGNT]-[EHW]-[DEFHLMNWY]-Y |
| | 70 | E-[ALMSTV]-D-P-[IM]-[DEGNT]-[EHW]-[DEFHLMWY]-Y |
| | 80 | E-[ALMSTV]-D-P-[IM]-[DEGNT]-[HW]-[DEFLWY]-Y |
| | 90 | E-[ALMSTV]-D-P-[IM]-[DEGNT]-[HW]-[DEFLWY]-Y |
| | 100 | E-[ALMSTV]-D-P-[IM]-[DEGNT]-[HW]-[DEFLWY]-Y |
| A3-010 | 5 | E-[ACFGHILMQSTV]-[DY]-P-[ILMV]-[ACDEFGHILMNPSTVWY]-[ACDEFGHIKLMNPQSTVWY]-[ACDEFGHIKLMNPQRSTVWY]-[MVY] |
| | 10 | E-[ACFHILMQSTV]-[DY]-P-[ILMV]-[ACDEFGHILMNPSTWY]-[ACDEFGHILMNPQSTVW]-[ACDEFGHIKLMNPQRSTVWY]-[VY] |
| | 20 | E-[ACHILMTV]-[DY]-P-[ILV]-[ACDEFGHILMNPSTY]-[ACEFGHILMNPQSTVW]-[ACDEFGHIKLMNPQRSTVWY]-Y |
| | 30 | E-[ACILMTV]-[DY]-P-[ILV]-[DEFGHILMNSTY]-[ACEGHILMNPQSTVW]-[ACDEFGHIKLMNPQRSTVWY]-Y |
| | 40 | E-[AIMV]-D-P-[ILV]-[DEFGHIMNST]-[ACEGHILMNPSTVW]-[ACDEFGHILMNPQRSTVWY]-Y |
| | 50 | E-[IMV]-D-P-[ILV]-[DEFGHMNST]-[ACEGHILNPSTVW]-[ACDEFGHLMNPQRSTWY]-Y |
| | 60 | E-[MV]-D-P-[ILV]-[DEFGHNST]-[ACEHILNSTVW]-[ACDEFHLMNPQSTWY]-Y |
| | 70 | E-[MV]-D-P-[ILV]-[DEGHT]-[AEHILNSTV]-[ACDEFLMNQSWY]-Y |
| | 80 | E-[MV]-D-P-[ILV]-[DEGT]-[AHILNSV]-[CDEFLMNSWY]-Y |
| | 90 | E-[MV]-D-P-I-[DEGT]-[AHILSV]-[CDEFLSWY]-Y |
| | 100 | E-[MV]-D-P-I-[DEG]-[AHILSV]-[DEFLY]-Y |

**Supplementary table 5.** Selected candidate targets of $TCR_{a3a}$ as predicted by DMS of the MAGE-A3 peptide and motif querying of the UniProtKB database

| Peptide code | UniProtKB entry | Protein | Protein full name | Species | Sequence | Activation threshold (%) | Predicted affinity HLA-A*0101 (nM) |
|---|---|---|---|---|---|---|---|
| MAGE | P43357 | MAGA3_HUMAN | Melanoma-associated antigen 3 | Human | EVDPIGHLY | 100 | 11.43 (SB) |
| Titin | Q8WZ42 | TITIN_HUMAN | Titin | Human | ESDPIVAQY | 80 | 8.07 (SB) |
| RVL-1 | P43360 | MAGA6_HUMAN | Melanoma-associated antigen 6 | Human | EVDPIGHVY | 70 | 39.73 (SB) |
| RVL-2 | Q07075 | AMPE_HUMAN | Glutamyl aminopeptidase | Human | ELYPMIEEY | 50 | 6402.84 (WB) |
| RVL-3 | Q6P6B7 | ANR16_HUMAN | Ankyrin repeat domain-containing protein 16 | Human | EGDPLILQY | 50 | 492.12 (SB) |
| RVL-4 | Q13740 | CD166_HUMAN | CD166 | Human | EMDPVTQLY | 50 | 5.68 (SB) |
| RVL-5 | P26010 | ITB7_HUMAN | Integrin beta-7 | Human | EGYPVDLYY | 50 | 5535.99 (WB) |
| RVL-6 | Q16621 | NFE2_HUMAN | Transcription factor NF-E2 45 kDa subunit | Human | EMYPVEYPY | 50 | 2707.36 (WB) |
| RVL-7 | Q2HR95 | AN_HHV8P | Shutoff alkaline exonuclease (SOX) | Kaposi's sarcoma-associated herpesvirus | ECDPIYAAY | 50 | 114.78 (SB) |
| RVL-8 | Q5VT25-2 | MRCKA_HUMAN | Serine/threonine-protein kinase MRCK alpha, isoform 2 | Human | ETDPVENTY | 50 | 7.26 (SB) |
| RVL-9 | Q96M61 | MAGBI_HUMAN | Melanoma-associated antigen B18 | Human | EVDPIRHYY | 10 | 18.05 (SB) |
| RVL-10 | Q1HVE7 | AN_EBVA8 | Shutoff alkaline exonuclease (SOX) | Epstein-Barr virus | EFDPIYPSY | 10 | 5267.99 (WB) |
| RVL-11 | Q18BH3 | RIMP_PEPD6 | Ribosome maturation factor RimP | C. difficile | EKDPIKENY | N/A | 20510.08 |

**Supplementary table 6.** Sequences of DNA oligonucleotides utilized in this study

| Purpose | Oligo name | Sequence 5'-3' | Annealing temp (°C) | Figure(s) |
|---|---|---|---|---|
| GFP PCR | RVL-63 | gatcagaagacgaagaagcatggtgagcaagggcgaggagc | 72 | S1e |
| | RVL-64 | gacttgaagacctcttaccttacttgtacagctcgtccatgccgag | | |
| CD8α-P2A-CD8β PCR | RVL-50 | cgtacaggatccgtcatggccttaccagtgaccgc | 72 | S1e |
| | RVL-51 | tgctcaacgcgtgtactgtcgactaagatacattgatgagtttgg | | |
| CD4 PCR | RVL-101 | gatgttgggacggcgataatgga | 65 | S3c |
| | RVL-102 | tgtggagctgaggcaacaaagaa | | |
| Labeling of cDNA 3' ends | TSO | AAGCAGTGGTATCAACGCAGAGTGAATrGrG+G | n/a | S5a, S7a |
| Jurkat TCRα template-switching RT-PCR | ISPCR | aagcagtggtatcaacgcagagt | 64 | S5a |
| | RVL-70 | tctcagctggtacacggcag | | |
| Jurkat TCRβ template-switching RT-PCR | ISPCR | aagcagtggtatcaacgcagagt | 62 | S7a |
| | RVL-68 | agatctctgcttctgatggctc | | |
| Jurkat TCRα RT-PCR | RVL-69 | cagtcggtgacccagcttgg | 62 | S5e |
| | RVL-70 | tctcagctggtacacggcag | | |
| NFAT-GFP HDR template generation (AAVS1) | RVL-119 | tgctttctctgaccagcattctctc | 72 | S4a |
| | RVL-120 | agagcagagccaggaaccc | | |
| AAVS1 PCR 1 | RVL-137 | cacctactcagacaatgcgatgc | 60 | S4e |
| | RVL-138 | gaactctgccctctaacgctgc | | |
| AAVS1 PCR 2 | RVL-139 | ctgggataccccgaagagtgagt | 65 | S4e |
| | RVL-140 | ccgcctggaaaggttagaggaaa | | |
| TnT-TCR PCR | RVL-71 | gtgcccttctcttttgttggcaag | 61.5 | 1e |
| | RVL-72 | gattcaggcagaggttgggagtt | | |
| TnT-TCR RT-PCR A (amplicon for deep seq.) | RVL-144 | gaggagaaccctggacctatg | 60 | S9a |
| | RVL-145 | ggaacaccttgttcaggtcctc | | |
| TnT-TCR RT-PCR B | RVL-67c | ttctttgctcatgctcacagagg | 62 | S9b |
| | RVL-68 | agatctctgcttctgatggctc | | |
| Plasmid library PCR (amplicon for deep seq.) | RVL-144 | gaggagaaccctggacctatg | 66 | S11c, S11d, S15b |
| | RVL-154 | ctagagacccccagccttacc | | |
| Primary T cell RT-PCR 1 | TRBV5-1_fwd | atcctggtaccaacagaccccaggacag | 68 | 5b |
| | TRACex3_rev1 | gtcatgagcagattaaacccggccac | | |
| Primary T cell RT-PCR 2 | TRAV21_fwd | atggaaaccctcttgggcctg | 62.3 | 5b |
| | TRACex3_rev2 | aaacccggccactttcag | | |
| TCRαβ HDR template generation (TnT cells) | RVL-127 | gcatgcctctgtgccaacag | 70 | 1b, S11b, S15a |
| | RVL-128 | ttttatctgtcatggccgtgaccg | | |
| TCRβα HDR template generation (1° T cells) | RVL-166 | ctgcctttactctgccagagttatattgc | 62 | 5a |
| | RVL-167 | gacatcattgaccagagctctggg | | |

| | | | | |
|---|---|---|---|---|
| Nicking mutagenesis $TCR_{A3}$ DMS | A3_NNK1 | ggccctttatctttgcnnkagcagcccgaatatggcg | n/a | S11 |
| | A3_NNK2 | ggccctttatctttgtgctnnkagcccgaatatggcggatgaaca | | |
| | A3_NNK3 | ggccctttatctttgtgctagcnnkccgaatatggcggatgaaca | | |
| | A3_NNK4 | ggccctttatctttgtgctagcagcnnkaatatggcggatgaacagtac | | |
| | A3_NNK5 | ggccctttatctttgtgctagcagcccgnnkatggcggatgaacagtac | | |
| | A3_NNK6 | tctttgcgccagcagcccgaatnnkgcggatgaacagtac | | |
| | A3_NNK7 | cgccagcagcccgaatatgnnkgatgaacagtacttcg | | |
| | A3_NNK8 | gcagcccgaatatggcgnnkgaacagtacttcg | | |
| | A3_NNK9 | ccgaatatggcggatnnkcagtactttgggccgggcaccagg | | |
| | A3_NNK10 | gaatatggcggatgaannktacttcgggcccggcaccaggctc | | |
| | A3_NNK11 | atatggcggatgaacagnnkttcgggcccggcaccaggctc | | |
| Nicking mutagenesis $TCR_{DMF4}$ DMS | DMF4_NNK1 | cagacatctgtgtacttctgtnnkatcagtgaggtaggggttggg | n/a | S11 |
| | DMF4_NNK2 | atctgtgtacttctgtgccnnkagtgaggtaggggttgggc | | |
| | DMF4_NNK3 | gtgtacttctgtgccatcnnkgaggtaggggttgggcag | | |
| | DMF4_NNK4 | tacttctgtgccatcagtnnkgtaggggttgggcagccc | | |
| | DMF4_NNK5 | ctgtgccatcagtgagnnkggggttgggcagcccc | | |
| | DMF4_NNK6 | gtgccatcagtgaggtannkgttgggcagccccagca | | |
| | DMF4_NNK7 | ccatcagtgaggtagggnnkgggcagccccagcattt | | |
| | DMF4_NNK8 | tgtgccatcagtgaggtaggggttnnkcagccccagcatt | | |
| | DMF4_NNK9 | cagtgaggtaggggttgggnnkccccagcattttggtgatg | | |
| | DMF4_NNK10 | gaggtaggggttgggcagnnkcagcattttggtgatggg | | |
| | DMF4_NNK11 | gtaggggttgggcagcccnnkcattttggtgatgggact | | |
| | DMF4_NNK12 | gggttgggcagccccagnnktttggtgatgggactcg | | |
| Nicking mutagenesis (2nd strand synthesis) | RVL-128 | ttttatctgtcatggccgtgaccg | n/a | S11 |
| Combinatorial $TCR_{A3}$ library OE-PCR | A3_fwd_ultramer | atcctggtaccaacagaccccaggacagggccttcagttcctctttgaatactt cagtgagacacagagaaacaaaggaaacttccctggtcgattctcagggc gccagttctctaactctcgctctgagatgaatgtgagcaccttggagct | 70 | S15a |
| | A3_rev_ultramer | taggctctcctagagacccccagccttacctgtgaccgtgagcctggtgccc ggcccgaarwaywgyycryynrcvnbrytvnbrcyrcyngsgcaaagata aagggccgagtcccccagctccaaggtgctcacattcatctcag | | |
| Combinatorial $TCR_{A3}$ library flanking PCR | A3_flank_fwd | atcctggtaccaacagaccccaggacag | 62 | S15a |
| | A3_flank_rev | taggctctcctagagacccccagccttacc | | |

rG = riboguanosine; +G = locked nucleic acid guanine; r = a, g; y = c, t; s = g, c; w = a, t; k = g, t; m = a, c; b = c, g, t; d = a, g, t; h = a, c, t; v = a, c, g; n = any base.

**Supplementary table 7.** Sequences of custom Alt-R crRNAs (IDT) utilized in this study

| Name | Target cells | Sequence (5' - 3') |
|---|---|---|
| CCR5 gRNA | Pre-TnT | tgacatcaattattatacat |
| GFP gRNA | Pre-TnT | caactacaagacccgcgccg |
| AAVS1 gRNA | Pre-TnT | ggggccactagggacaggat |
| CD4 gRNA | Pre-TnT | gcactgaggggctactacca |
| TRAC gRNA (Jurkat) | Pre-TnT | cagggttctggatatctgt |
| Fas gRNA | Pre-TnT | ttggaaggcctgcatcatga |
| CDR3β gRNA | TnT cells | tcgacctgttcggctaacta |
| TRAC gRNA | Primary T cells | agagtctctcagctggtaca |
| TRBC1/2 gRNA | Primary T cells | ggagaatgacgagtggaccc |
